# Enhanced dynamic covalent chemistry for the controlled release of small molecules and biologics from a nanofibrous peptide hydrogel platform

**DOI:** 10.1101/2024.05.21.595134

**Authors:** Brett H. Pogostin, Samuel X. Wu, Michael J. Swierczynski, Christopher Pennington, Si-Yang Li, Dilrasbonu Vohidova, Erin H. Seeley, Anushka Agrawal, Chaoyang Tang, Jacob Cabler, Arghadip Dey, Omid Veiseh, Eric L. Nuermberger, Zachary T. Ball, Jeffrey D. Hartgerink, Kevin J. McHugh

## Abstract

Maintaining safe and potent pharmaceutical drug levels is often challenging. Multidomain peptides (MDPs) assemble into supramolecular hydrogels with a well-defined, highly porous nanostructure that makes them attractive for drug delivery, yet their ability to extend release is typically limited by rapid drug diffusion. To overcome this challenge, we developed self-assembling boronate ester release (SABER) MDPs capable of engaging in dynamic covalent bonding with payloads containing boronic acids (BAs). As examples, we demonstrate that SABER hydrogels can prolong the release of five BA-containing small-molecule drugs as well as BA-modified insulin and antibodies. Pharmacokinetic studies revealed that SABER hydrogels extended the therapeutic effect of ganfeborole from days to weeks, preventing *Mycobacterium tuberculosis* growth better than repeated oral administration in an infection model. Similarly, SABER hydrogels extended insulin activity, maintaining normoglycemia for six days in diabetic mice after a single injection. These results suggest that SABER hydrogels present broad potential for clinical translation.

## Main

Many drugs have narrow therapeutic windows or short half-lives, necessitating frequent administration and exacerbating the potential for side effects when administered improperly.^1^ Drug delivery systems can enhance therapeutic efficacy by retaining drug concentrations within the therapeutic window for extended periods of time, alleviating medication-related side effects associated with peak-to-trough variations, maximizing drug exposure, and improving patient adherence to medication schedules.^2, 3^ Supramolecular peptide hydrogels are a particularly promising class of biomaterials for drug delivery that have been evaluated in preclinical and clinical settings. They offer favorable biocompatibility and readily degrade into non-toxic products.^4–6^

Multidomain peptides (MDPs) are one class of supramolecular peptide hydrogels with a programmable and well-defined nanostructure consisting of a highly porous nanofibrous network.^7^ MDPs self-assemble into bilayered β-sheet-rich nanofibers with a cross-section of 2 nm by 6 nm while the length of individual nanofibers extends for microns.^8^ Their nanostructure allows the formation of physically crosslinked hydrogels at low concentration (1% by weight) that can transiently disassemble under shear stress,^9, 10^ allowing them to be easily injected through a needle yet form a hydrogel bolus in vivo that will remain in a fixed location for weeks.^11, 12^ Because of these excellent properties, MDPs have been investigated for the delivery of diverse payloads, such as carbohydrates,^13^ proteins,^14^ and small molecules,^15^ yet challenges associated with insufficient release lifetime remain due to the small size of these payloads relative to the mesh size of the hydrogel.^16^

Affinity-based release modalities have been employed to prolong drug release from hydrogels, including complementary electrostatic interactions, guest-host binding, hydrogen bonding, hydrophobic interactions, and dynamic covalent interactions.^17–21^ Dynamic covalent interactions are stronger than electrostatic or hydrophobic interactions and form in dynamic equilibrium under physiologically relevant conditions, allowing for drug release.^22^ Dynamic covalent bonding between diol and boronic acid (BAs) motifs forms boronate esters and has been leveraged in the design of drug delivery systems.^23–25^ While 1,2-dihydroxyphenyl derivatives (catechols) are the most commonly used motif in BA dynamic covalent chemistry,^26–28^ they have limited affinity for BAs and experience oxidative decomposition that can accelerate drug release, reduce payload stability, and lead to crosslinked byproducts.^29–33^ Furthermore, competition for BA binding between catechols and glucose can result in release rates that fluctuate greatly with blood glucose levels.^34^

Herein, we overcome the limitations of previous boronate ester-mediated release systems by functionalizing self-assembling peptides with 4-nitrocatechol or salicylhydroxamic acid (SHA) motifs to enhance boronate ester formation and stability in order to extend the duration of drug release.^26, 29, 35, 36^ We developed a small library of <u>s</u>elf-<u>a</u>ssembling <u>b</u>oronate <u>e</u>ster release (SABER) MDP hydrogels capable of engaging in dynamic covalent bonding with BAs to extend the release of BA-containing small molecules (BACSMs), a small-molecule drug modified with a BA, and BA-modified biologics without interfering with the peptide’s ability to rapidly self-assemble into shear-thinning and self-healing nanofibrous hydrogels (**Fig. 1a**). The ability to release multiple classes of drugs from SABER hydrogels—in combination with the simplicity and modularity of the platform—make it well-suited for clinical translation.

**Fig 1.**
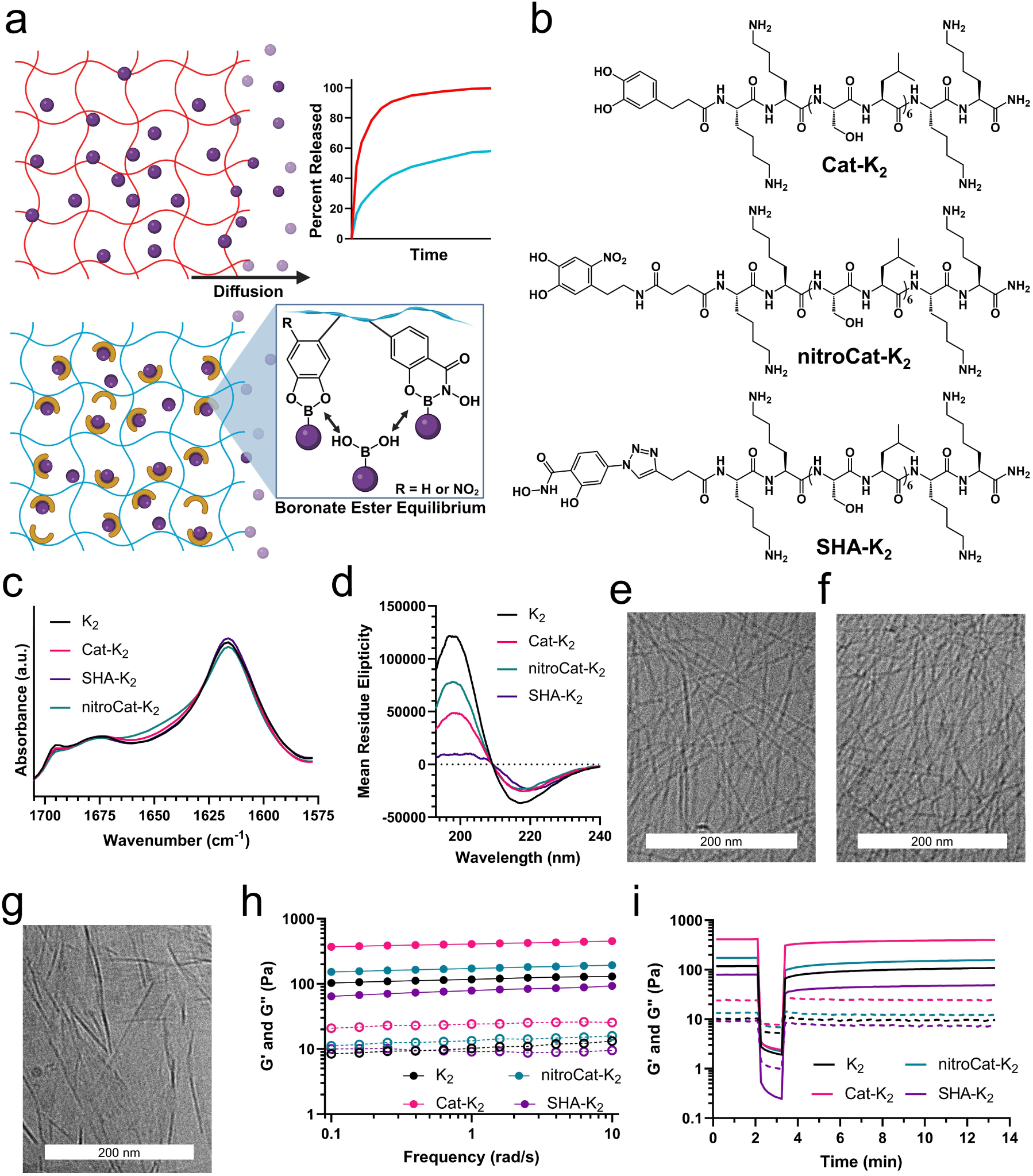
Characterization of SABER hydrogels. a) A schematic illustrating how small molecules rapidly diffuse from peptide hydrogels due to their large mesh size (red). We hypothesized that by adding dynamic covalent attachment moieties to the hydrogel, release can be slowed resulting in prolonged delivery (blue). b) Chemical structures of the three SABER peptides synthesized. c) FTIR of the three SABER peptides shows that they assemble into antiparallel β-sheets. d) CD spectra of unmodified K2 and SABER MDPs are consistent with β-sheet secondary structure. e-g) *Cryo*-TEM images of (e) Cat-K_2_, (f) nitroCat-K_2_, and (g) SHA-K2 show that all peptides self-assemble into nanofibers. The scale bar = 200 nm. h) Oscillatory rheology frequency sweeps show that the three SABER peptides at 10 mg/mL form hydrogels with similar moduli to unmodified K2. i) Shear recovery rheology experiments where all four peptides were subjected to 200% strain for 1 min, demonstrating that all the hydrogels are shear thinning and rapidly self-healing.

### Design and development of SABER nanofibrous hydrogels

Motivated by the shortcomings of catechol-BA bonding, we aimed to identify chemical motifs that (1) engaged in dynamic covalent bonding with BAs, (2) resisted oxidation, and (3) exhibited an equilibrium shifted more towards the bonded (boronate ester) state to increase drug retention in MDP hydrogels. Longitudinal UV-Vis analysis showed that, at ambient temperature in pH 7.4 phosphate-buffered saline (PBS), SHA and 4-nitrodopamine demonstrated excellent stability over 15 d while a model catechol, dopamine, rapidly degraded (**Extended Data Fig. 1a-c**). Investigations into the boronate ester association constants between catechol, SHA, and 4-nitrodopamine and several BACSMs of interest indicated that SHA formed the strongest dynamic covalent interactions, exceeding the catechol association constants by 6.5- to 9.6-fold depending on the BACSM (**Extended Data Fig. 1d**).

Based on the oxidation and bonding results, we synthesized the MDP Ac-K_2_(SL)_6_-K_2_-Am (referred to as K_2_) functionalized with catechol (Cat-K_2_), 4-nitrocatechol (nitroCat-K_2_), or SHA (SHA-K_2_) motifs to yield a set of three SABER peptides (**Fig. 1b; Extended Data Fig. 1e-h**). These modified MDPs retained the ability to self-assemble into nanofibrous hydrogels, as assessed by Fourier transform infrared (FTIR) spectroscopy, circular dichroism (CD), rheology, and cryogenic transmission electron microscopy (*cryo*-TEM). The FTIR spectra of all four peptides displayed peaks at 1618 and 1695 cm^-1^, consistent with anti-parallel β-sheet secondary structure (**Fig. 1c**),^8, 10^ which was confirmed by the presence of a minimum ca. 216-220 nm in the CD spectra of all the peptides (**Fig. 1d**). Cat-K_2_ and nitroCat-K_2_ peptides formed long thin nanofibers as seen by *cryo*-TEM, whereas SHA-K_2_ nanofibers appeared to cluster and form sheet-like bundles of nanofibers (**Fig. 1e-g**). Rheological testing confirmed that all four peptides formed hydrogels, as indicated by storage moduli (G’) that exceed the loss moduli (G”) at all oscillatory frequencies tested (**Fig. 1h**). When subjected to 200% strain, G” exceeded G’ for all gels, suggesting liquid-like flow under high shear. Hydrogel character rapidly recovered once the shear force was reduced, suggesting these materials reform as hydrogels after injection (**Fig. 1i**).

To determine if dynamic covalent bonding to BAs could prolong the release of BACSMs from peptide hydrogels, we evaluated the *in vitro* release of five different small molecules from the nanofibrous SABER hydrogels. Using a model BA-modified fluorophore, we found that SABER MDPs drastically reduced the release rate of the fluorophore compared to K_2_ only when modified with a BA (**Extended Data Fig. 2a & b**). Next, we sought to demonstrate the controlled *in vitro* release of the FDA-approved BACSMs bortezomib (BTZ) and ixazomib (multiple myeloma therapeutics), the FDA-approved antifungal agent tavaborole, and ganfeborole (GFB, a tuberculosis treatment in phase II clinical trials).^25, 37^ In the cases of BTZ and ixaxomib, which have similar boronic acid motifs, unmodified K_2_ and Cat-K_2_ hydrogels released >82.3% and >57.1% of each drug in the first 2 h, respectively. In contrast, SHA-K_2_ and nitroCat-K_2_ hydrogels only released 15-25% of each drug in 2 h and 50-70% within 24 h (**Fig. 2a & b**). A reduction in the initial burst release was observed with Cat-K_2_ when releasing tavaborole, with 80.0 ± 4.7% of the cargo released in 24 h (**Fig. 2c**). SHA-K_2_ and nitroCat-K_2_, however, exhibited significantly lower tavaborole release at 24 h (49.1 ± 0.3% and 64.8 ± 0.9% released, respectively) than Cat-K_2_ and K_2_. For GFB, however, both K_2_ and nitroCat-K_2_ had similarly rapid release rates (**Fig. 2d**), consistent with the low equilibrium constant between GFB and 4-nitrodopamine shown in Extended Data Fig. 1d. The release of GFB from Cat-K_2_ could not be quantified due to the rapid oxidative deboronation of the compound (**Extended Data Fig. 2c-f**), suggesting that the catechol facilitates oxidative destruction of the drug.^31, 38^ Only SHA-K_2_ was able to significantly extend the release of GFB compared to K_2_, releasing less drug in 24 h (48.7 ± 2.5%) than K_2_ released in 30 min (59.0 ± 1.1%). Changing the drug loading by varying the drug-to-peptide molar ratio had a limited effect on the peptide secondary structure (**Extended Data Fig. 2g**) and release rate below a drug-to-peptide ratio of 1:1 (**Fig. 2e & f**; additional discussion in the Supporting Information).

**Fig 2.**
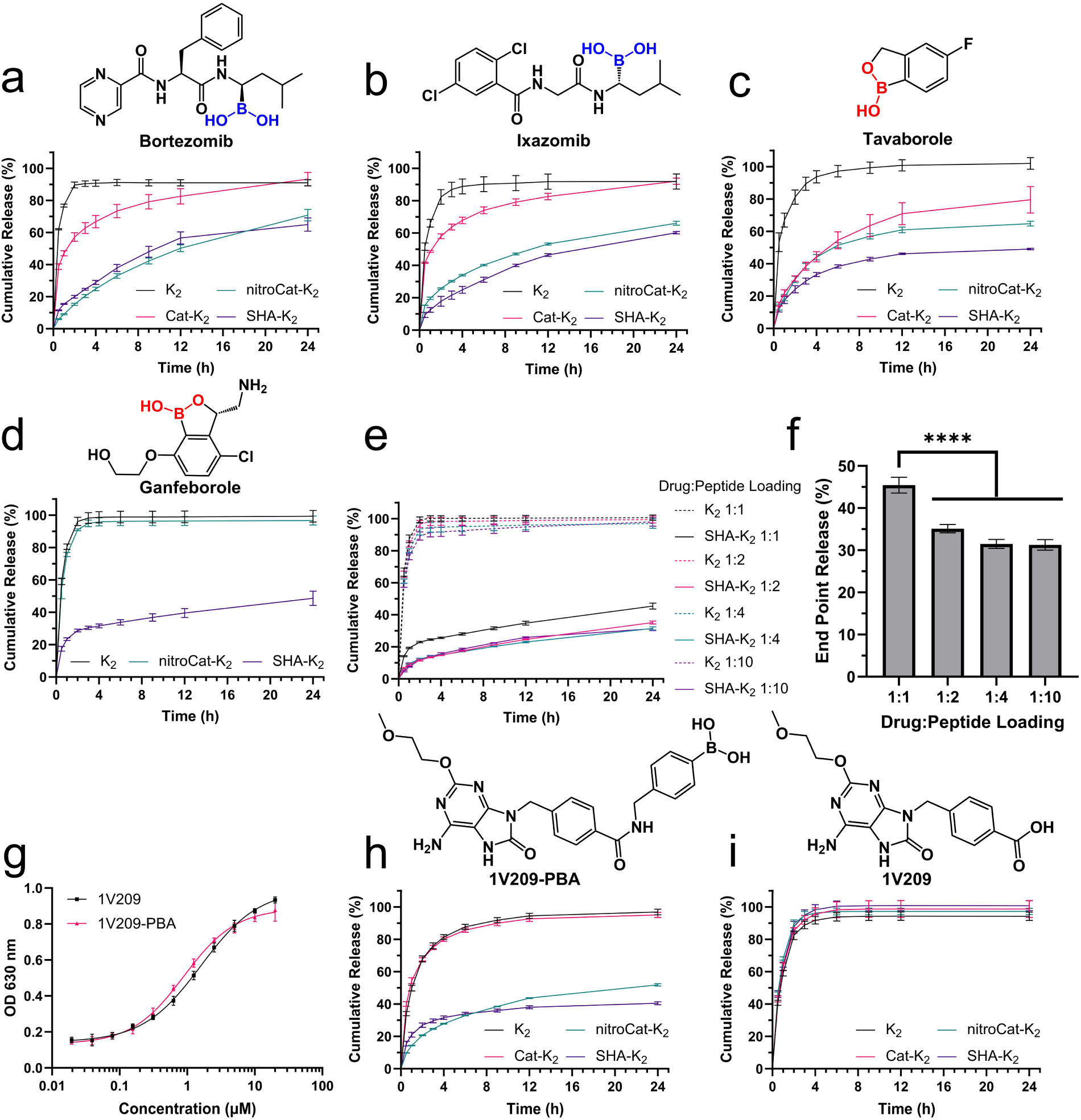
*In vitro* release of boronic acid-containing small molecule drugs. *In vitro* release o*f* (a) bortezomib and (b) ixazomib loaded at 0.5 mg/mL shows that SABER hydrogels prolong the delivery of chemically distinct drugs with BAs (shown in blue) compared to unmodified K_2_. c) Release of tavaborole from hydrogels at 0.25 mg/mL showed that SABER hydrogels are compatible with benzoxaborole-containing drugs (indicated in red). d) SHA-K_2_ was the only peptide able to significantly slow the release of GFB loaded at 0.5 mg/mL *in vitro*. e-f) Changing the drug-to-peptide molar ratio does not significantly impact the release rate until approaching a 1:1 ratio. The release of GFB from unmodified K2 did not vary as a function of drug loading. g) Adding PBA to 1V209 did not affect the biological activity as determined by a colorimetric TLR7 activity assay performed using the HEK-Blue mTLR7 reporter cell line. *In vitro* release of (h) 1V209-PBA and (i) 1V209 show that the PBA motif is necessary for SABER hydrogels to extend the release of the drug. The release of 1V209-PBA from Cat-K_2_ was identical to the unmodified K2 hydrogel. Data points indicate mean ± 1 SD (n=3).

Although BAs have become important moieties in medicinal chemistry,^25^ most compounds used in the clinic do not contain a BA. To expand the potential applications of the SABER platform, we examined the release of a BA-containing analog of a small-molecule drug that does not have a BA. We modified 1V209, a toll-like receptor 7 (TLR7) agonist, with a phenylboronic acid (PBA) motif, yielding 1V209-PBA, and found that PBA modification did not impact its ability to activate TLR7 relative to the parent drug in a TLR7 reporter cell line (**Fig. 2g**). *In vitro* release studies revealed 1V209-PBA exhibited dramatically slower release from nitroCat-K_2_ and SHA-K_2_ than Cat-K_2_ and K_2_ hydrogels (**Fig. 2h**). In the absence of the PBA modification, no altered release kinetics were observed (**Fig. 2i**). These results underscore the functional importance of the novel modifications of MDPs with nitroCat or SHA, which succeed at extending drug release from hydrogels when the catechol modification fails, and provide a workflow to make other small-molecule drugs compatible with the SABER delivery platform.

### *In vivo* release of BACSM therapeutics from SABER hydrogels

BTZ was the first BACSM to achieve FDA approval and is the most studied BACSM in the context of drug delivery.^39^ To evaluate the controlled release of BTZ, from SABER hydrogels *in vivo*, BALB/c mice were subcutaneously injected with 700 ng of BTZ alone or the same mass of BTZ loaded in 50 μL of MDP hydrogel. Mice receiving BTZ loaded in nitroCat-K_2_ and SHA-K_2_ experienced a significantly lower C_max_ (30.4 ± 2.0 ng/mL and 17.0 ± 4.3 ng/mL, respectively) and the C_max_ was observed significantly later (34 h) compared to BTZ alone or in K_2_ (**Fig. 3a & b**). A high C_max_ after BTZ administration in patients has been linked to toxicity;^40^ therefore, the more than 6-fold reduction in BTZ C_max_ could reduce side effects. We then performed a dose de-escalation study of BTZ alone to determine the dose that results in a C_max_ that matches that of 700 ng delivered from SABER hydrogels (**Extended Data Fig. 3a**). We found that a dose of 175 ng of BTZ alone yielded a C_max_ of 21.2 ± 3.1 ng/mL, which is statistically similar to the C_max_ observed in mice dosed with a 5-fold higher BTZ dose (700 ng) released from nitroCat-K_2_ and SHA-K_2_ (**Extended Data Fig. 3b**). At this peak-matched dose of BTZ, mice dosed with BTZ loaded in nitroCat-K_2_ and SHA-K_2_ hydrogels had a 2.9-fold and 1.6-fold higher drug exposure (AUC), respectively, and a 3-fold higher circulating BTZ concentration 336 h following administration than the peak matched dose without a hydrogel (**Fig. 3c**).

**Fig 3.**
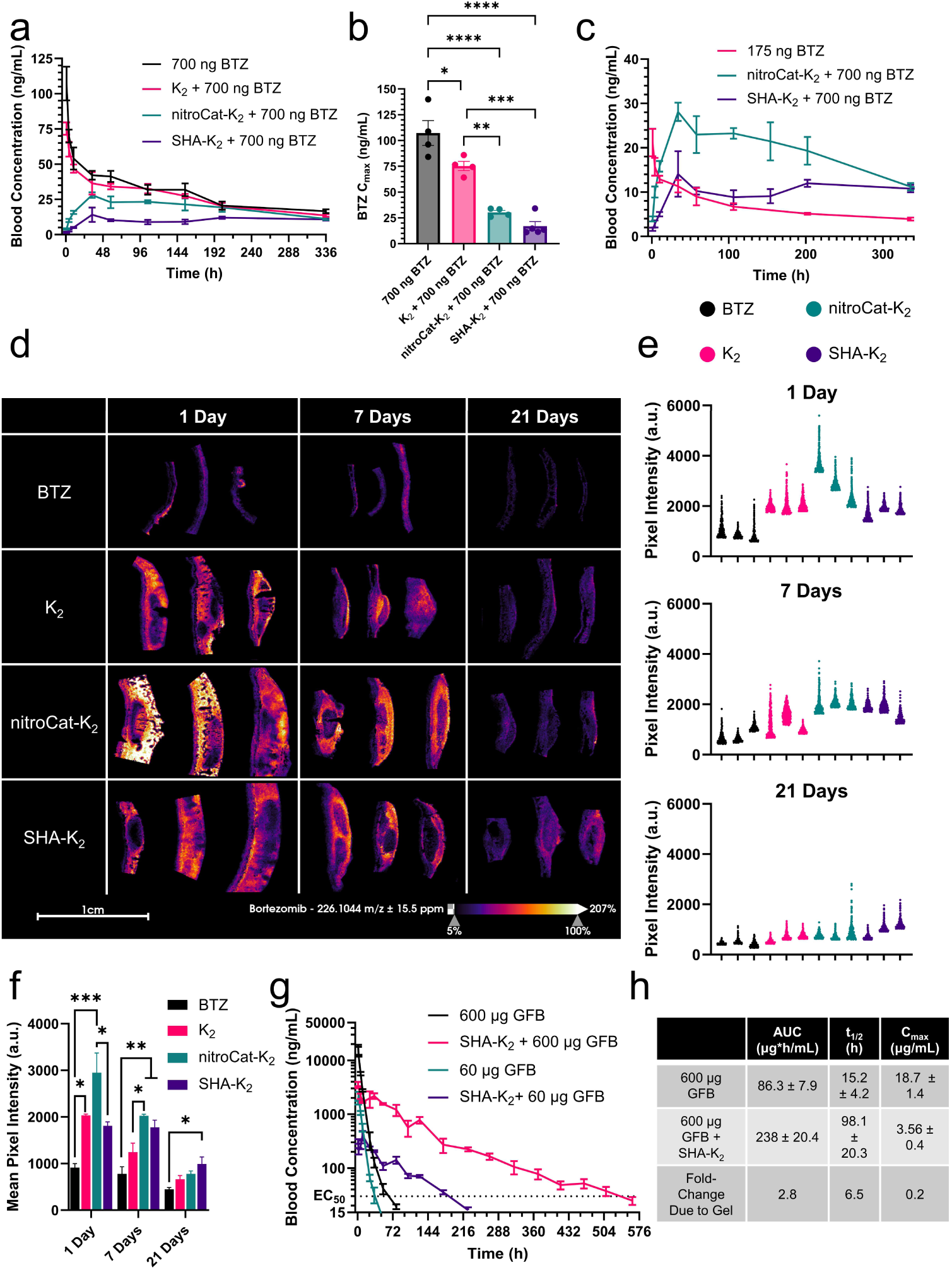
SABER peptides prolong the systemic release and local retention of BACSMs *in vivo*. a) Pharmacokinetic profile of 700 ng of BTZ administered subcutaneously without a hydrogel or loaded in 10 mg/mL K2, nitroCat-K_2_, or SHA-K_2_ (n=4-5). b) Delivering BTZ from nitroCat-K_2_ and SHA-K_2_ hydrogels significantly reduces the maximum circulating concentration (Cmax) of the drug compared to BTZ alone or BTZ loaded in K2. c) BTZ without a hydrogel must be administered at a 5-fold lower dose (175 ng) to match the Cmax achieved from delivering 700 ng of BTZ from nitroCat-K_2_ and SHA-K_2_. d) Mass spectrometry imaging analysis of BTZ at the injection site shows that MDP hydrogels retain higher local concentrations of BTZ. Darker spots in tissues are hydrogels, which suppress BTZ ionization. e) Pixel intensities over the 1 mm^2^ of tissue with the highest BTZ concentration at (top) 1 d, (middle) 7 d, and (bottom) 21 d. f) At early time points, the mean pixel intensity in nitroCat-K_2_ is the highest of all the groups, while SHA-K_2_ is the only group statistically superior to BTZ alone at 21 d. g) Pharmacokinetic profile of 600 (∼1:1 molar ratio of drug-to-peptide) and 60 µg (∼1:10 molar ratio of drug-to-peptide) of GFB administered alone or in a 20 mg/mL SHA-K_2_ hydrogel shows that the time above the EC50 is improved by increasing drug loading (n=4). h) Non-compartmental pharmacokinetic analysis of the groups dosed with 600 µg revealed that the SHA-K_2_ significantly improved the drug exposure (AUC) and half-life (t1/2) of GFB while reducing the Cmax. All data points are shown as mean ± SEM.

We then investigated the persistence of the drug at the site of injection, which could be beneficial for some applications, such as treating solid tumors.^41, 42^ Mass spectrometry imaging of tissue sections collected from the injection site showed that mice receiving BTZ loaded in MDPs have higher levels of signal attributable to BTZ (**Extended Data Fig. 3c & d**), compared to mice receiving the drug alone, at all time points (**Fig. 3d & Extended Data Fig. 4a**). We observed that at 1 and 7 d post-injection, the tissue surrounding nitroCat-K_2_ hydrogels had the highest BTZ concentrations, but at 21 d, only the drug levels surrounding the SHA-K_2_ injection sites were statistically superior to unformulated BTZ (**Fig. 3e & f**). Overlaying the BTZ signal with that of heme confirmed that the BTZ signal being measured is from the tissue rather than blood vessels (**Extended Data Fig. 4b**). These data suggest that the duration of local release can be controlled using different BA dynamic covalent bonding motifs.

To further demonstrate the compatibility of the SABER platform with multiple drugs and disease applications, we evaluated the *in vivo* release of the tuberculosis (TB) drug GFB. TB treatment typically requires months of daily oral drug dosing and is fraught with low patient adherence, which is as low as 27% among high-risk populations.^43, 44^ Improving patient adherence to TB treatment has motivated the field to identify long-acting injectable formulations.^3^ Consistent with the *in vitro* release results, mice receiving 75 μg of GFB in 50 μL of 10 mg/mL K_2_ and nitroCat-K_2_ gels were unable to prolong GFB release *in vivo* compared to the drug alone, whereas SHA-K_2_ gels maintained GFB blood concentrations above the EC_50_ of the drug for over 4-fold longer than other groups.^45^ (**Extended Data Fig. 5**). Since GFB is well tolerated in humans at high doses,^45^ we sought to extend the duration over the EC_50_ by increasing the peptide concentration, the volume of hydrogel administered, and the drug loading. Mice were subcutaneously injected with 200 µL of 20 mg/mL SHA-K_2_ hydrogels loaded with 600 µg and 60 µg of GFB. At both drug concentrations, loading GFB into SHA-K_2_ significantly prolonged the release of the drug compared to the drug alone and resulted in a 5.2-fold reduction in the C_max_ (**Fig. 3g & h**). Mice that received 600 µg of GFB in SHA-K_2_ had circulating GFB concentrations above the EC_50_ for over 508 h compared to under 79 h in mice that received the drug alone. Non-compartmental pharmacokinetic analysis of the 600 µg dose revealed that SHA-K_2_ improved the AUC by 2.8-fold and increased the circulating half-life (t_1/2_) of GFB by 6.5-fold (**Fig. 3h**).

### SABER peptide modularity and ability to improve TB treatment

To demonstrate that the SABER platform is compatible with various self-assembling peptides, we changed the base MDP in our SHA-modified SABER hydrogel from K_2_ to Ac-EE(SL)_6_EE-Am (E_2_), a negatively charged peptide (**Fig. 4a; Extended Data Fig. 6a & b**). The strongly cationic K_2_ MDP is known to cause an immune response,^14, 46^ which may be well-suited to cancer applications but makes it less ideal for conditions like TB that do not benefit from inflammation. Negatively charged MDPs have been found to cause minimal inflammation *in vivo*.^46^ SHA-modified E_2_ (SHA-E_2_) spontaneously self-assembled into an antiparallel β-sheet secondary structure, like the base peptide, as indicated by FTIR (**Fig. 4b**) and CD (**Fig. 4c**). SHA-E_2_ also formed hydrogels with similar rheological properties to E_2_ (**Extended Data Fig. 6c**). Both E_2_ and SHA-E_2_ hydrogels were shear thinning and self-healing and recovered 90% and 88% of their initial G’ within 10 min, respectively, after being subjected to 200% strain (**Fig. 4d**). *Cryo*-TEM images revealed the presence of thin twisted ribbon nanofibers (**Fig. 4e**).

**Fig 4.**
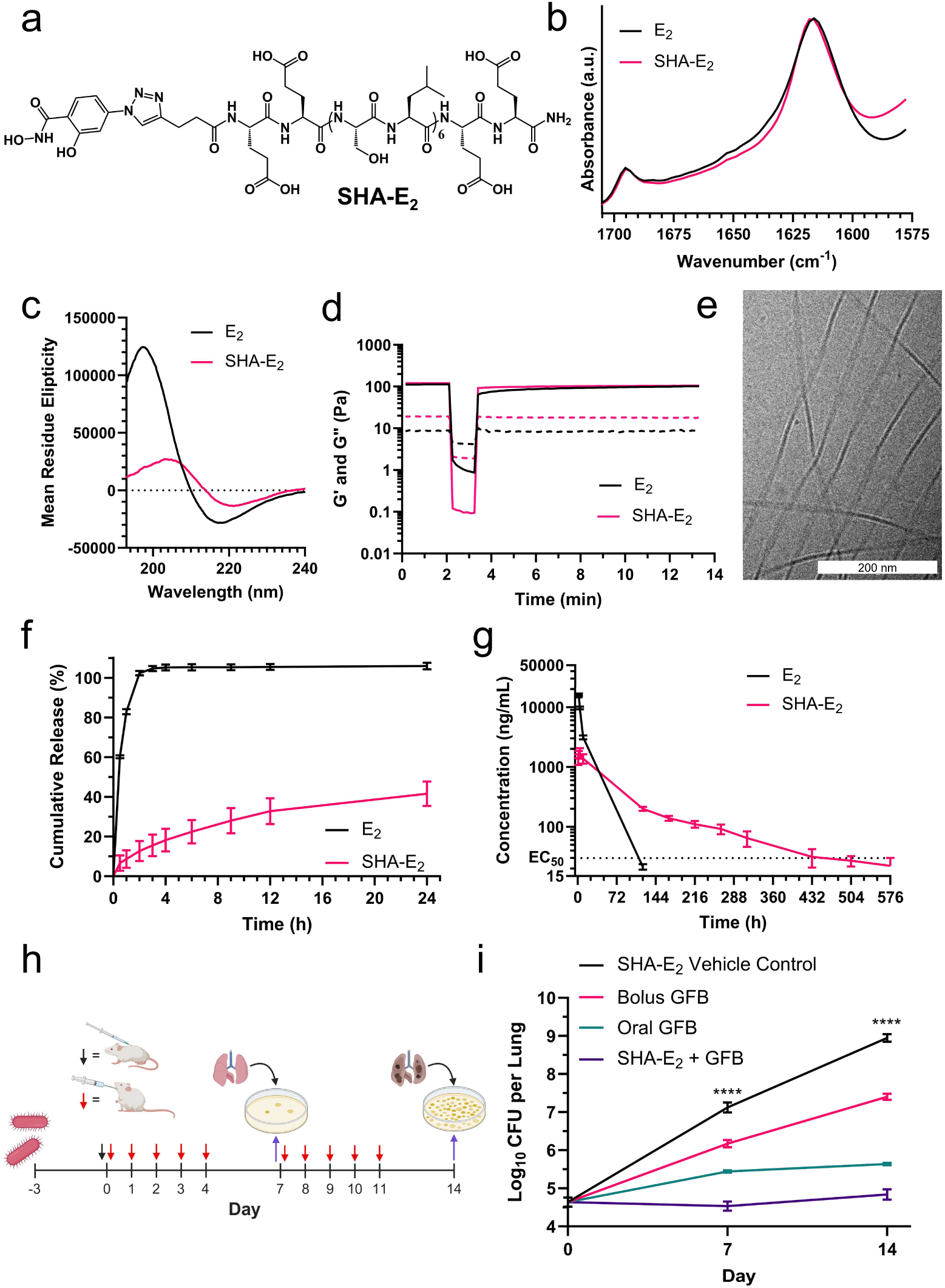
Characterization of negatively charged SABER hydrogels and their use to treat a murine model of acute TB. a) Chemical structure of SHA-E_2_. b) FTIR of SHA-E_2_ shows that the antiparallel β-sheet secondary structure of the peptide remains similar to unmodified E_2_. c) CD spectra of both E_2_ and SHA-E_2_ indicate the presence of β-sheet self-assembly. d) Rheological testing shows that both E_2_ and SHA-E_2_ hydrogels are shear thinning and recover their viscoelastic properties within 10 min after being subjected to 200% strain for 1 min. e) C*ryo*-TEM images of SHA-E_2_ peptides in solution showing self-assembled nanofibers. Scale bar = 200 nm. f) SHA-E_2_ significantly prolongs the release of GFB compared to E_2_ *in vitro* when loaded at 0.5 mg/mL. Data points indicate mean ± 1 SD (n=3). g) Pharmacokinetic profile after dosing 600 µg of GFB shows that SHA-E_2_ prolongs the release of the drug *in vivo* compared to unmodified E2. Data points indicate mean (n=5) ± SEM. h) A schematic illustrating the experimental design for the mouse model of acute TB. i) The CFU counts from the lungs of mice with TB over 14 days show that a single injection of 600 μg GFB in SHA-E_2_ outperforms the same dose injected without the gel or the same total mass of drug given over 10 oral doses. CFU data are presented as the mean ± 1 SD (n=4).

*In vitro*, SHA-E_2_ significantly reduced the release of GFB compared to the E_2_ control and released 41.6 ± 6.2% of the total drug in 24 h (**Fig. 4f**), which is statistically similar to the cumulative release measured from SHA-K_2_ under the same conditions (**Extended Data Fig. 6d**). An *in vivo* pharmacokinetic study using 200 µL of 20 mg/mL SHA-E_2_ loaded with 600 µg of GFB similarly showed that SHA-E_2_ prolongs drug release compared to the unmodified E_2_ hydrogel. This SHA-E_2_ formulation maintained a circulating GFB concentration above the EC_50_ for more than 400 h and enhanced the t_1/2_ of the compound by over 10-fold compared to GFB alone (**Fig 4g & Extended Data Fig. 6e**).

After demonstrating that SHA-E_2_ could extend the release of GFB *in vivo*, we investigated the use of this platform to treat acute TB. Three days after aerosol infection with *M. tuberculosis*, mice were treated with 600 μg of GFB delivered as a single subcutaneous bolus in PBS or SHA-E_2_. Since GFB is given orally to treat TB in clinical trials, an additional group was treated orally with the same total mass of drug administered across 10 doses over the course of 14 d. Mice were euthanized 7 and 14 d after the start of the treatment to quantify the colony-forming units (CFU) of *M. tuberculosis* in the lungs (**Fig. 4h**). Mice given the SHA-E_2_ hydrogel without any drug rapidly showed signs of an uncontrolled bacterial infection over 14 d (**Fig. 4i**). Giving the drug without hydrogel as a single injection reduced the CFU by 0.95 and 1.55 log_10_ at 7 and 14 d, respectively. The effect of the drug was improved when given in 10 oral doses, which resulted in a reduction of 1.67 log_10_ CFU after 7 d and 3.32 log_10_ CFU after 14 d compared to the vehicle control. A single injection of GFB loaded in SHA-E_2_ significantly enhanced the efficacy of the treatment compared to all groups and resulted in a 2.58 log_10_ and 4.12 log_10_ reduction in CFU compared to the vehicle control at 7 and 14 d, respectively. This nearly 10-fold reduction in CFU compared to mice that received oral dosing suggests that using SABER hydrogels could simultaneously reduce the necessary dosing frequency and enhance the efficacy of GFB compared to the oral dosing strategy used in the clinic.

### Prolonged local delivery of a BA-modified IgG from SABER

Building on the success of SABER to deliver BACSMs, we adapted the system to biological payloads. Although biotherapeutics, such as monoclonal antibodies, have risen in popularity owing to their high specificity and potency, their clinical translation can be limited by poor pharmacokinetics and/or adverse effects.^47^ We hypothesized that modifying antibodies with a BA-motif would enable SABER hydrogels to retain antibodies for long periods of time *in vivo* (**Fig. 5a**). Thus, a model fluorescent rabbit IgG was modified with either a low number of PBAs (IgG Low, 2.4 PBAs per antibody) or a high number of PBAs (IgG High, 11.4 PBAs per antibody). Fluorescence recovery after photobleaching experiments demonstrated that both SHA-K_2_ and SHA-E_2_ reduce the diffusion of BA-modified IgG within the gels (**Fig 5b & c**). We additionally found that the number of PBA motifs per IgG had a minimal impact on IgG diffusion within the range tested (**Extended Data Fig. 7a & b**, additional discussion in the Supplementary Information).

**Fig 5.**
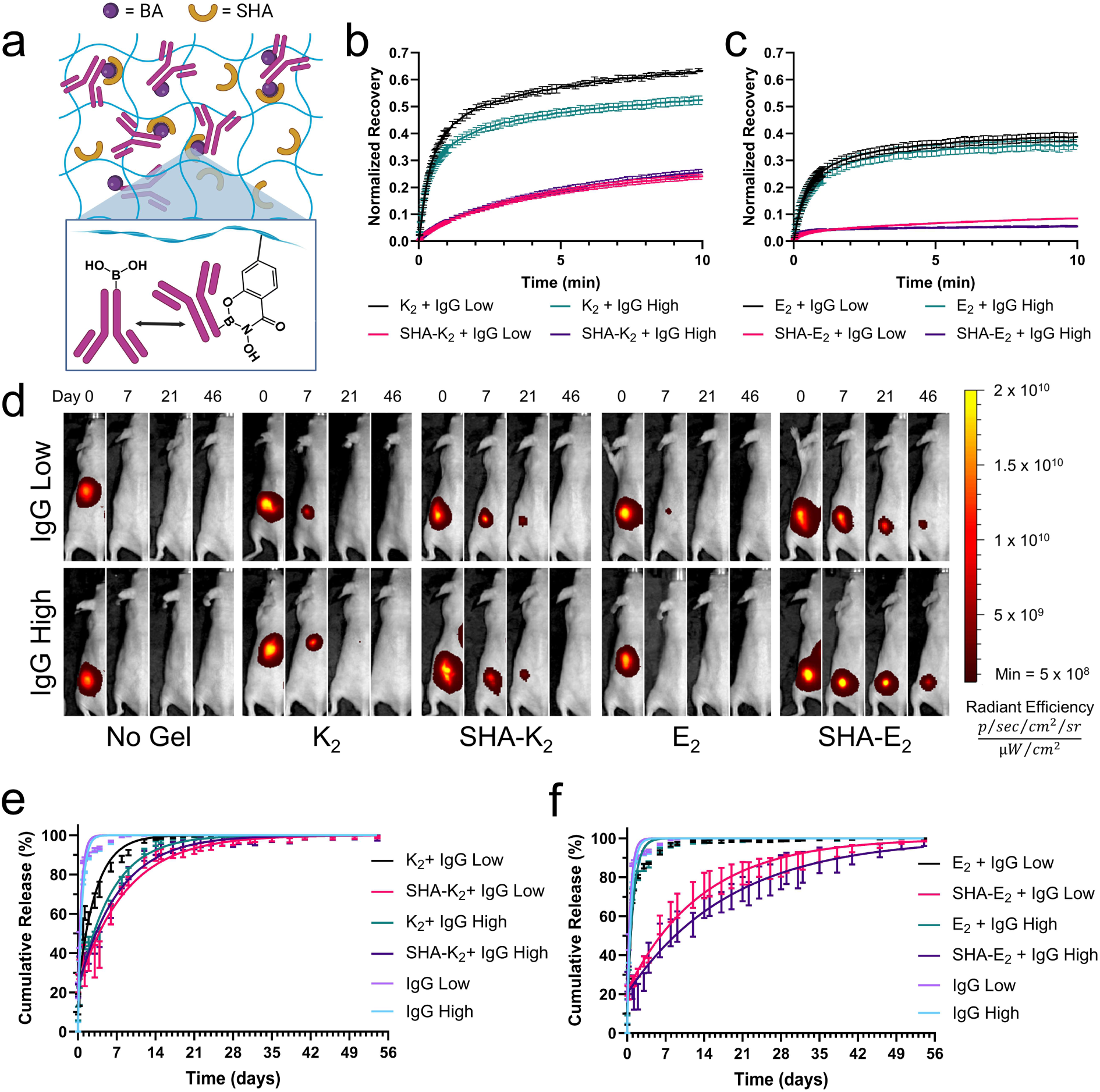
Local delivery of a BA-labeled model IgG antibody from SABER hydrogels. a) A schematic illustrating dynamic covalent bonds forming between BA motifs on labeled IgG molecules and SABER hydrogels to slow down transport within the hydrogel. b & c) The normalized fluorescence intensity recovery after photobleaching a region of (b) SHA-K_2_ or (c) SHA-E_2_ hydrogels loaded with fluorescently labeled IgG with 2.4 PBAs per IgG (IgG Low) and 11.4 PBAs per IgG (IgG High). SHA modification of MDPs slowed down the transport of IgG within the hydrogel matrix compared to unmodified MDPs. Data are presented as the mean ± 1 SD (n=3). d) Representative *in vivo* fluorescence images of mice subcutaneously injected with negatively and positively charged SABER hydrogels loaded with fluorescent IgG Low and IgG High at different time points. e & f) Quantification of changes in fluorescence intensity at the injection site over time in mice receiving PBA-labeled IgG shows that SHA-K_2_ and SHA-E_2_ hydrogels prolong the release of the antibodies to more than 28 and 56 days, respectively. Data points indicate mean (n=4) ± SEM.

To elucidate the ability of SABER hydrogels to slow the release of BA-modified antibodies *in vivo*, fluorescence imaging was used to quantify the local retention of fluorescently labeled IgG High and IgG Low injected subcutaneously into mice (**Fig. 5d**). Without a hydrogel, IgG Low and IgG High injections were rapidly cleared after administration. K_2_ moderately improved the retention of IgG despite the absence of dynamic covalent bonding (**Fig. 5e**). In the case of IgG Low, using SHA-K_2_ instead of K_2_ more than doubled the t_1/2_ of the payload from 2.2 to 5.9 d (Extended Data Fig. 7c). All groups using K_2_ as a base peptide, including SHA-K_2_ gels, had minimal (< 5%) IgG remaining after 4-weeks. In comparison, SHA-E_2_ extends the retention of PBA-modified IgG out to 8 weeks (**Fig. 5f**). Using SHA-E_2_ to deliver IgG Low resulted in an almost 24-fold improvement in the t_1/2_ of the antibody alone from 0.4 to 9.5 d and an almost 14-fold improvement compared to unmodified E_2_ (**Extended Data Fig. 7c**). Increasing labeling from 2.4 to 11.4 PBAs per antibody led to modest but statistically significant improvements in payload retention (**Extended Data Fig. 7c**). These data suggest that minimal levels of BA labeling are required to significantly improve local antibody retention and that SHA-E_2_ is superior to SHA-K_2_ in releasing PBA-labelled antibodies.

### SABER enables days-long glycemic control in diabetic mice

We explored the flexibility of this platform by using it to deliver another therapeutically relevant protein, insulin. There is a persistent need for insulin delivery systems due to the high clinical burden of basal insulin therapy.^3, 48^ Many groups have investigated using boronate ester chemistry to create glucose-responsive insulin delivery systems.^49–53^ This responsiveness is desirable for meal-related insulin dosing but is undesirable for basal insulin delivery, which aims to provide long-term normalization of insulin levels. The boronate ester association constant between SHA and PBA is >4 x 10^3^-fold higher than the previously reported value for the glucose-PBA interaction (Extended Data Fig. 1d).^27^ As such, we hypothesized that SHA-E_2_ could be formulated as a glucose-insensitive delivery system for PBA-modified insulin (insulin-PBA; **Extended Data Fig. 8a & b**) to induce prolonged normoglycemia in diabetic mice.

*In vitro* release assays in varying concentrations of glucose (0, 100, and 250 mg/dL) showed that PBA modification was necessary to delay release (**Extended Data Fig. 8c**) and that SHA-E_2_ significantly slowed the release of insulin-PBA at all glucose concentrations compared to E_2_ (**Fig. 6a**). A modest, yet statistically significant, increase in insulin release was observed at 250 mg/dL glucose compared to 0 mg/dL (**Fig. 6b**). Healthy blood glucose concentrations range from 80-180 mg/dL and a blood glucose level over 240 mg/dL can require urgent intervention; therefore, SABER hydrogels are unlikely to be meaningfully impacted by physiologically relevant glucose levels.^54^

**Fig 6.**
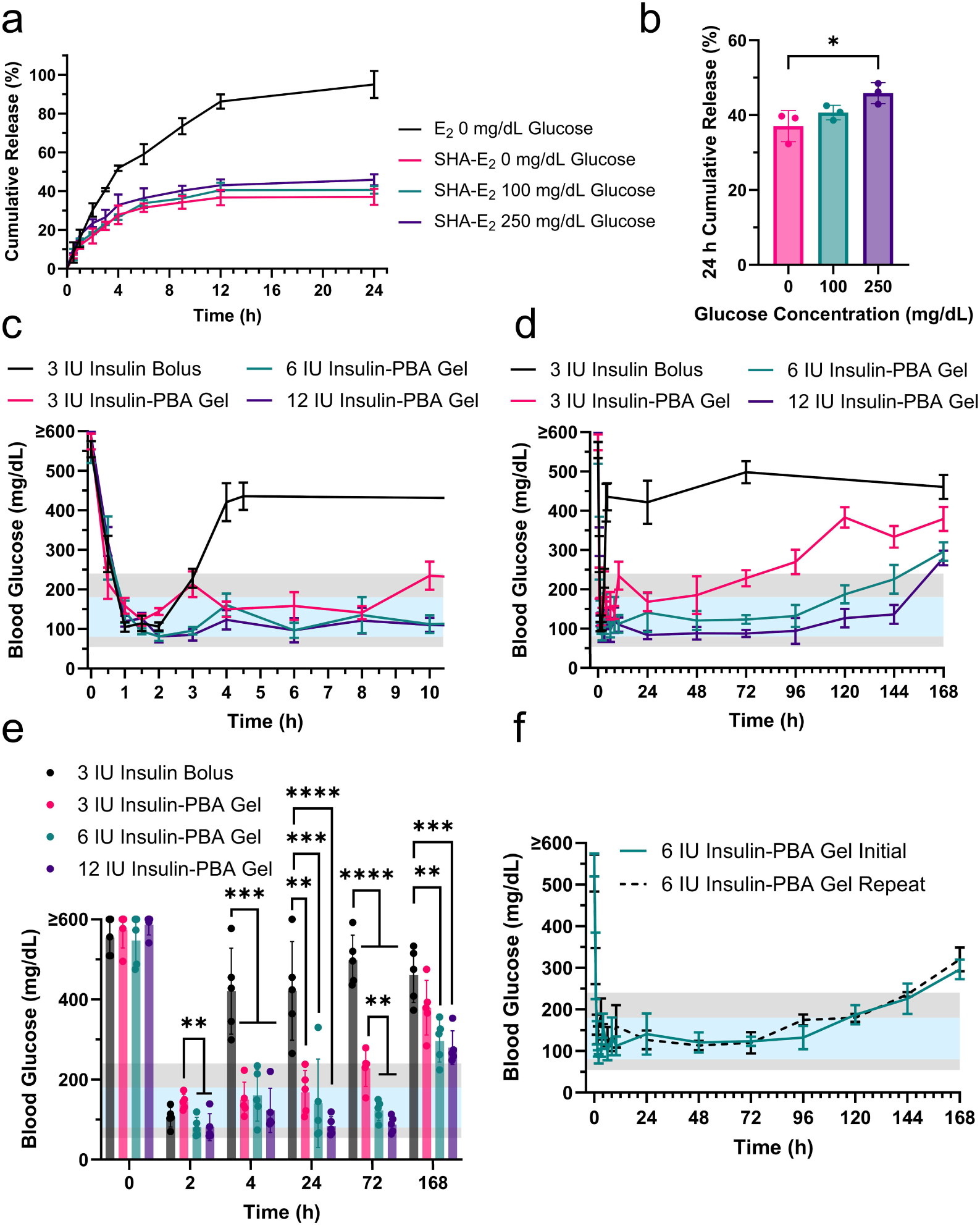
Delivery of PBA-modified insulin from SHA-E_2_ maintains normoglycemia for up to 144 h in a mouse model of type 1 diabetes. a) *In vitro* release of 0.5 mg/mL of insulin-PBA loaded in E_2_ or SHA-E_2_ in media with 0, 100, or 250 mg/dL of glucose. Data plotted as the mean ± 1 SD (n=3). b) The cumulative 24 h *in vitro* release of insulin-PBA from SHA-E_2_ reveals that the hydrogel is not likely glucose responsive over healthy blood glucose concentrations. Error bars represent ± 1 SD from the mean (n=3). c) Diabetic mice treated with a subcutaneous bolus of unmodified insulin without a hydrogel return to hyperglycemic blood glucose levels less than 4 h after injection while mice administered 3, 6, or 12 IU of insulin-PBA within SHA-E_2_ remain below 240 mg/dL. d) The duration over which insulin-PBA delivered in SHA-E_2_ can maintain normoglycemia in diabetic mice increases with the loading of insulin-PBA in the gel. e) At 4, 24, and 72 h after administration, SHA-E_2_ loaded with insulin-PBA results in significantly lower blood glucose levels than a single bolus of 3 IU of insulin. At 168 h, both 6 and 12 IU insulin-PBA gels are statistically superior to the 3 IU bolus control even though mice are hyperglycemic. f) Retreating mice with 6 IU of insulin-PBA loaded in SHA-E_2_ 6 weeks after the initial treatment yielded statistically similar control of blood glucose levels. The blue region of all blood glucose graphs represents healthy blood glucose levels in humans (80-180 mg/dL), the grey regions represent mild hyperglycemia (> 240 mg/dL) and mild hypoglycemia (54-80 mg/dL), and white regions represent critically low or high blood glucose levels. All blood glucose data points show mean (n=5) ± SEM.

We then tested whether a single injection of insulin-PBA loaded in an SHA-E_2_ hydrogel could maintain normoglycemia in diabetic mice for multiple days. Mice received a subcutaneous injection of SHA-E_2_ hydrogels loaded with 3, 6, or 12 IU of insulin-PBA or 3 IU of unmodified insulin without a hydrogel. Immediately after administration, the blood glucose concentration in all mice rapidly fell below 150 mg/dL; however, mice that received 3 IU of insulin without a hydrogel became hyperglycemic again within 4 h **(Fig. 6c**). The average blood glucose level in mice receiving 3 IU of insulin-PBA in the SHA-E_2_ remained below 240 mg/dL for 72 h (**Fig. 6d**), effectively controlling blood glucose levels 18-fold longer than 3 IU of unmodified insulin.

Mice that received gels loaded with 6 IU remained below 180 mg/mL for 96 h and below 240 mg/dL for 144 h, while mice administered 12 IU remained below 180 mg/dL for 144 h (Fig. 6d). The 6 and 12 IU insulin-PBA gel treatment groups achieved statically significantly lower blood glucose levels than the insulin only control between 4 and 168 h and outperformed the 3 IU gel group consistently after 72 h (**Fig. 6e**). The blood glucose levels of mice treated with a second dose of 6 IU of insulin-PBA in an SHA-E_2_ hydrogel were not statistically significantly different from those obtained with the first dose at all time points measured (**Fig. 6f & Extended Data Fig. 8d**). These data establish that insulin-PBA loaded in SHA-E_2_ can be used to achieve normoglycemia for up to 144 h in a single injection, a 36-fold improvement over the maximum tolerated single dose of unmodified insulin without a hydrogel.

## Conclusions

Modifying MDPs such that they can dynamically form boronate esters with therapeutics significantly enhances their capacity to control the release of BA-containing payloads while leaving their nanostructure and rheological properties intact. Using nitrocatechol and SHA, novel BA dynamic covalent association motifs for drug delivery applications, we show that SABER hydrogels can be used to enhance the pharmacokinetics and/or distribution of BA-containing compounds. The SABER drug delivery platform is compatible with a wide array of different modalities including synthetically BA-modified drugs, whether those be small molecules or proteins, to greatly extend the duration of drug activity *in vivo*. The enhanced dynamic covalent bonding strength afforded by SHA modification allows for the extended release of BA-containing small molecules and biologics for weeks instead of days *in vivo*. In this paper, we demonstrate the utility of this system in several clinically relevant models with different classes of drugs, including the ability to better control TB with a single injection than with repeated oral dosing and the ability to achieve normoglycemia in diabetic mice for 144 h after a single injection. Taken together, these results establish the SABER hydrogel platform as a highly flexible drug delivery system that may have the potential to improve the treatment of a wide variety of different diseases.

## Acknowledgements

The authors would like to acknowledge Ulf Olsson and the Olsson laboratory for their hospitality at Lund University and Crispin Hetherington for capturing *cryo*-TEM images. The authors recognize Felipe Lerner for assistance synthesizing an SHA starting material compatible with solid-phase peptide synthesis, Gabriel Saenz for his assistance with peptide synthesis, Hossam Kandry for guidance with pharmacokinetic analyses, and Alloysius J. Budi Utama for help developing a protocol for FRAP. The authors would like to further acknowledge Neeraja Dharmaraj, Nourhan Hussein, Simon Young, and Andrew Sikora for their support and advice throughout the project. BHP received funding from the NSF Graduate Student Research Fellowship program and the National Cancer Institute F99/K00 program (award number: F99CA284262). Mass spectrometry imaging was performed in the UT Austin Mass Spectrometry Imaging Facility supported by Cancer Prevention and Research Institute of Texas award RP190617. This project was supported in part with NIH grants R35GM143101, R01DE021798, R01DE030140, and R61-AI-161809. We acknowledge support from the Welch Foundation (Research Grant C-1680), the National Science Foundation (CHE-2203948), and the Cancer Prevention and Research Institute of Texas (RR190056).

## Methods

### Supplementary Methods

Methodology for chemical synthesis, UV-Vis, binding assays, TLR7 activation assays, and fluorescence recovery after photobleaching is included in the Supplementary Information.

### Hydrogel Preparation

MDP hydrogels were prepared by dissolving the MDP at 2X the final desired peptide concentration (20 mg/mL or 40 mg/mL) in MilliQ water. The desired BACSM or boronic acid-labeled biologic was dissolved at twice the final concentration in 2X Hank’s balanced salt solution (HBSS). The pH of the 2X HBSS solution was adjusted to pH 7.5-8.5 to assist in the dissolution of BACSMs. These two stock solutions were then mixed in a 1:1 ratio to yield a hydrogel with 1X HBSS, and the final 1X peptide and payload concentrations (10 mg/mL or 20 mg/mL). The pH of the gel was adjusted to pH 7-8 as necessary with microliter additions of NaOH or HCl and the gel was vortexed to ensure homogeneity.

### Fourier Transform Infrared Spectroscopy (FTIR)

Attenuated total reflectance FTIR was performed using a Nicolet^TM^ iS20 FTIR spectrometer (Thermo Fisher Scientific). Before use, a stream of nitrogen was used to purge the instrument to avoid signals from atmospheric molecular vibrations. Hydrogel samples at a peptide concentration of 10 mg/mL were plated (10 μL) and allowed to dry until all water had evaporated. Absorbance spectra were collected as an accumulation of 30 measurements. The amide I FTIR peak was visualized by background subtraction and plotting the area normalized absorbance from 1575-1705 cm^-1^.

### Circular Dichroism (CD) Spectroscopy

Circular dichroism spectra were obtained on a Jasco J-810 spectropolarimeter (Easton, MD). Hydrogel samples (10 µL) with or without drug were loaded into a 0.1 mm quartz cuvette and analyzed at ambient temperature. All spectra were obtained as an average of 5 accumulations between 190 and 250 nm. Millidegrees (mdeg) of rotation were converted to mean residual ellipticity (MRE) for ease of comparison using the equation [θ] = θ × 10^6^ /(*c* × *l* × *nr*), wherein *c* is concentration (μmol/L), *l* is cuvette pathlength (mm), *nr* is the number of amino acid residues, and θ is the ellipticity reading in mdeg.

### Rheology

Oscillatory rheology was performed on an AR-G2 rheometer (TA Instruments, New Castle, DE). Rheology measurements were collected from 75 μL hydrogel samples using a stainless steel 12 mm parallel plate geometry. After plating, samples were allowed to equilibrate at 1 rad/s and 1% strain for 30 min. After equilibration, a frequency sweep was performed over the range of 0.1-10 rad/s at a constant 1% strain. Subsequently, shear recovery experiments were conducted by equilibrating the hydrogels at 1 rad/s and 1% strain for 2 min and then sheared for 1 min at 200% strain. The recovery of G’ and G” was subsequently monitored for 10 min at the starting conditions.

### Cryogenic Transmission Electron Microscopy (*cryo*-TEM)

Hydrogels were prepared as previously described at 10 mg/mL in 1X HBSS buffer. Before plunging, samples were diluted to 1 mg/mL in fresh MQ water. Samples (5 μL) were placed on 200 mesh lacey carbon grids and then were plunged into liquid ethane with a Leica EM GP automatic plunge freezer (Leica Microsystems, Wetzlar, Germany). The plunging chamber was set to a temperature of 20 °C and a humidity of 90%. Samples were analyzed at the National Center for High-Resolution Electron Microscopy at Lund University, Sweden, on a JEM-2200FS (JOEL, Tokyo, Japan) transmission electron microscopy instrument equipped an F416.0 camera (TVIPS, Gauting, Germany) using an accelerator voltage of 200 kV. Serial EM in a low-dose mode were used to acquire zero-loss images.

### *In Vitro* Release Assays

#### Plate Reader Release Assay

Hydrogels at a final peptide concentration of 10 mg/mL were loaded with 1.5 mM fluorescein or FITC-PBA were prepared as previously described and allowed to equilibrate overnight at ambient temperature in the dark. To start the experiment, 50 μL of each hydrogel was plated in triplicate on to a custom 3D-printed 48-well plate designed previously for plate reader release assays,^14^ and the gels were allowed to recover for 10 min. The gels were then covered gently with 600 μL of 1X PBS, sealed with a transparent plate sticker to prevent evaporation, and loaded into a microplate reader (Tecan, Zurich, Switzerland). The microplate reader was set to heat the plate to 37 °C while orbitally shaking it at 142 rpm, and it was set to read the fluorescence (490/525 nm Ex/Em) of the release supernatant every 45 min. The percentage of payload released was calculated by dividing the fluorescence intensity at each time point to the total fluorescence of a solution of 0.115 mM of fluorescein or FITC-PBA, representing the concentration of cargo at 100% release.

#### BACSM UPLC Release Assay

All BACSMs were purchased from Ambeed except GFB, which was purchased from MedChemExpress (Monmouth Junction, NJ). Hydrogels at a final peptide concentration of 10 mg/mL were loaded with 500 µg/mL BACSM, were prepared as previously described earlier in the methods and equilibrated in the dark at ambient temperature overnight. Tavaborole (250 µg/mL), 1V209 (50 µg/mL), and 1V209-PBA (50 µg/mL) gels were prepared with lower concentrations due to solubility limitations in the hydrogels. Additionally, hydrogels with 1V209 and 1V209-PBA were prepared with 25% DMSO to improve the solubility of these hydrophobic small molecules. Gels (50 µL) were then plated in triplicate on a custom 3D-printed 48-well plate as described in previous publications. The gels were then allowed to recover for 10 min before 450 µL of 1X PBS was gently added to each well to start the experiment. The plate was then covered with a 96-well plate cover sticker to prevent evaporation and orbitally agitated at 100 rpm at 37 °C. At the desired sampling time points, 400 µL of the release media was removed and refreshed. The amount of BACSM in the removed release medium was then assessed by UPLC using a Poroshell C18 (Agilent Technologies, Santa Clara, CA) UPLC column and adjusting for the volume remaining in the well after each sampling (Supplementary Information Table S1). The percentage of released BACSM was determined by dividing the cumulative mass of drug released by the total loaded in the hydrogel.

#### Insulin Release Assay

Unmodified insulin and insulin-PBA were loaded at 500 µg/mL in hydrogels containing 10 mg/mL peptide as described above using 1X PBS instead of HBSS to avoid the addition of glucose. Each hydrogel (50 µL) was then pipetted into low protein bind Eppendorf tubes (Hamburg, Germany) in triplicate and centrifuged to settle the gel in the bottom of the tube. After the gels equilibrated for 10 min, 450 µL of 1X PBS 0, 100, and 250 mg/dL of glucose was gently pipetted on top of each gel and incubated at 37 °C. At the desired sampling time point, 200 uL of release media was removed and replaced, and the concentration of insulin and insulin-PBA in the release medium was quantified by UPLC using a BEH C14 column (Waters Corp.) accounting for the volume remaining in the tube after each sampling (Supplementary Information Table S1). The percentage of released Insulin and insulin-PBA was determined by dividing the cumulative mass of the protein released by the total loaded in the hydrogel.

### Pharmacokinetic Assays

#### Bortezomib

All animal work was performed in compliance with an IACUC-approved protocol. Hydrogels loaded with 700 ng of BTZ were prepared as previously described at a final peptide concentration of 10 mg/mL. Female BALB/c mice (16-18 g) were injected subcutaneously with 50 μL of drug-loaded hydrogel or BTZ dissolved in HBSS. Blood (10 μL) was collected using an untreated Safe-T-Fill™ plastic hematocrit capillaries (RAM Scientific, Austell, GA) and spotted on a Whatman 903 Proteinsaver Card (Cytiva, Marlborough, MA). Blood cards were dried overnight protected from light and then stored with desiccant at 4 °C for up to 1 week or -80 °C for up to one month. Extractions were performed following a previously published protocol for BTZ.^55^ The compound was extracted off the blood card by using a 1/8-inch hole puncher to remove the center of each blood spot, corresponding to 2.4 μL of blood. The blood spots were then submerged in 40 μL of methanol containing 1 ng/mL apatinib (APExBIO, Houston, TX) as the internal standard. Extractions proceeded for 1 h at ambient temperature on an orbital shaker set to 100 rpm. The extraction solution was then removed and diluted 1:2 with water and analyzed by LC-MS.

Multiple Reaction Monitoring (MRM) LC-MS analysis of BTZ was carried out on an Agilent 6470B Triple Quadrupole (QqQ) mass spectrometer using apatinib as an internal standard. The MS system was interfaced to an Agilent 1290 Infinity ii LC system through an Agilent Jet Spray (AJS) electrospray ionization (ESI) source that was operated in the positive mode. Separations were carried out using a Water’s ACUITY Premier HHS T3 100 mm x 2.1 mm ID, 1.8 um column that was operated at 0.4 mL/min. Mobile phase A (MPA) was 0.1% formic acid in water, and mobile phase B (MPB) was 0.1% formic acid in acetonitrile. Initial LC conditions were 20%B up to 80%B over 4.0 min. The column was flushed at 80%B for 2.0 min and then re-equilibrated at 20%B for 2 min prior to the next injection.

The AJS Source and MS data acquisition settings were optimized to meet the needed sensitivity requirements. Briefly, the AJS source conditions were as follows, Gas Temp: 320 °C, Gas Flow: 8 L/min, Nebulizer Gas Pressure: 25 psi, Sheath Gas Temp: 400 °C, Sheath Gas Flow: 11 L/min, Capillary Voltage: 3800 V. QqQ MRM data acquisitions settings were as follows, Cell Accelerator Voltage: 5 V, Dwell Time: 50 ms. The MRM ion transitions for bortezomib and apatinib are shown in the supplementary information (Supplementary Information Table S2)

#### Ganfeborole

Pharmacokinetic analysis of GFB was performed nearly identically to the procedure described above for bortezomib with minor alterations. Hydrogels loaded with 60 or 600 μg GFB were prepared as previously described at a final peptide concentration of 10 mg/mL or 20 mg/mL depending on the experiment. Female BALB/c mice (16-18 g) were injected subcutaneously with the drug loaded hydrogel or GFB dissolved in PBS. Blood (10 μL) was collected using an untreated plastic capillary and spotted on a Whatman 903 Proteinsaver Card. Blood cards were dried overnight protected from light and then stored with desiccant at 4 °C for up to 1 week or -80 °C for up to one month. The compound was extracted off the blood card by using a 1/8-inch hole puncher to remove the center of each blood spot, corresponding to 2.4 μL of blood. The punches were then submerged in 40 μL of 90:10 methanol:water containing 5 ng/mL apatinib as an internal standard. The submerged blood spots were extracted at 37 °C for 1 h while being shaken at 100 rpm. The extraction solution was then removed and diluted 1:2 with water and analyzed by LC-MS.

MRM LC-MS analysis of GFB was carried out on an Agilent 6470B QqQ mass spectrometer using apatinib as an internal standard. The MS system was interfaced to an Agilent 1290 Infinity ii LC system through an AJS ESI source that was operated in the positive mode. Separations were carried out using a Water’s ACUITY Premier HHS T3 100 mm x 2.1 mm ID, 1.8 um column that was operated at 0.4 mL/min. MPA was 0.1% formic acid in water, and MPB was 0.1% formic acid in acetonitrile. Initial LC conditions were 10%B up to 60%B over 5.0 min from 5.0 min to 5.5 min the gradient was increased to 80%B. The column was flushed at 80%B for 2.5 min and then re-equilibrated at 10%B for 3 min prior to the next injection.

The AJS Source and MS data acquisition settings were optimized to meet the needed sensitivity requirements. Briefly, the AJS source conditions were as follows, Gas Temp: 320 °C, Gas Flow: 5 L/min, Nebulizer Gas Pressure: 30 psi, Sheath Gas Temp: 400 °C, Sheath Gas Flow: 12 L/min, Capillary Voltage: 3800 V. QqQ MRM data acquisitions settings were as follows, Cell Accelerator Voltage: 5 V, Dwell Time: 100 ms, MS1 (Q1) resolution: Unit, MS2 (Q3) resolution: Wide. The MRM ion transitions for GFB and apatinib are shown are shown in the supplementary information (Supplementary Information Table S3) The data collected was corrected for dilutions during sample preparation and plotted in GraphPad Prism 10. Pharmacokinetic parameters were determined by noncompartmental analysis using Ubiquity in RStudio.^56^

### Mass Spectrometry Imaging

Female BALB/c mice (16-18 g) were subcutaneously injected with 700 ng of BTZ loaded in 50 µL of 1X HBSS, K_2_, nitroCat-K_2_, or SHA-K_2_. Hydrogels were prepared as previously described. Mice were then sacrificed 1, 7, and 21 d post administration and the tissue at the injection sites were collected. The tissue was cryopreserved with liquid nitrogen without the use of any fixative. Mouse skin was cross sectioned at 12 μm thickness using at Thermo NX50 cryostat (Epredia, Kalamazoo, MI) and collected onto standard plus slides. Optical images of the slides were acquired at 4800 dpi using an Epson Perfection V600 Photo flatbed document scanner (Epson US, Los Alamitos, CA). Sections were coated with 10 mg/mL α-cyano-4-hydroxycinnamic acid matrix in 70% ACN, 0.1% TFA using an HTX M5 Robotic Reagent Sprayer (HTX Technologies, LLC, Chapel Hill, NC) as follows: 4 passes, nozzle temperature of 75 °C, flow rate of 100 μL/min, track speed of 1200 mm/min, track spacing of 3 mm, an HH track pattern, and a nozzle height of 40 mm. Serial sections were collected for H&E staining and were digitized using a Hamamatsu NanoZoomerSQ Digital Slide Scanner (Hamamatsu Photonics, Bridgewater, NJ).

Mass spectrometry images were acquired at 50 μm resolution in positive ion mode using a Bruker timsTOF fleX QTOF mass spectrometer (Bruker Daltonics) over the *m/z* range 50-1000 with a summation of 700 shots per pixel. Instrument tuning was as follows: a Funnel 1 RF of 100.0 Vpp, a Funnel 2 RF of 150.0 Vpp, a Multipole RF of 200.0 Vpp, a Collision Energy of 5.0 eV, a Collision RF of 600.0 Vpp, a Quadrupole Ion Energy of 5.0 eV, a Transfer Time of 60.0 μs, and a Pre Pulse Storage of 6.0 μs. BTZ was detected in tissue as the in source generated fragment at m/z 226.09 that was confirmed through MALDI analysis of a standard.

Image files loaded into SCiLS Lab Pro 2023b (Bruker Daltonics) for visualization and analysis. Data were root mean square normalized. H&E images were annotated using Hamamatsu NDP.view2 software for regions of gels in the sections. These annotations were transferred to SCiLS to create ROIs corresponding to areas of gel and non-gel within each sample. Intensities of BTZ from each pixel outside the gels were exported to a .CSV file using the SCiLS Lab API for R. Dark spots in the mass spectrometry images are a result of the hydrogel suppressing the ionization of BTZ, thus only drug in the tissue outside the hydrogels could be quantified (Extended Data Fig. 3d & 4a). To account for the different sizes of tissue collected, the pixel intensities over the 1 mm^2^ (400 pixels) of tissue with the highest BTZ signal were analyzed for statical comparisons in GraphPad Prism 10.

### *In Vivo* Fluorescence Animal Imaging IgG Release Assay

Sterile 10 mg/mL MDP hydrogels containing 2 mg/mL of AZ647-labeled IgG-PBA were prepared one day before the assay and stored at 4 °C overnight protected from light. SKH1-Elite mice obtained from Charles River Laboratories (Wilmington, MA) and tissue background autofluorescence was quantified at Ex/Em 640/700 nm for each mouse using an *In Vivo* Imaging System (IVIS) small animal imager (PerkinElmer, Waltham, MA). Background fluorescence intensity was subtracted from all subsequent images. Mice were subcutaneously injected bilaterally in the flank with 50 µL of sample or control and imaged longitudinally. The percentage of material released from the injection site was calculated by drawing equally sized region-of-interest rectangles around the injection sites and dividing the observed total radiant efficiency in that region by the maximum total radiant efficiency measured for that injection on the first day of the experiment. The data were fit to first-order exponential equation in GraphPad Prism 10 using a least-squares regression to model the release from the injection site. The error for parameters extracted from the model are reported as 95% confidence intervals.

### Mouse Model of Acute TB

*M, tuberculosis* H37Rv was grown in 7H9 broth supplemented with 10% oleic acid, albumin, dextrose, catalase ([OADC] Difco Laboratories, Detroit, MI) and 0.05% Tween 80 (Sigma-Aldrich, St. Louis, MO) before infection. A log phase growth (acute infection) model infection was used for this experiment. In brief, 6-week-old female BALB/c mice (Charles River Laboratories) were infected with a log-phase culture of *M. tuberculosis* (optical density at 600 nm of approximately 1.0) using an inhalation exposure system (Glas-Col, Terre Haute, IN), aiming to implant approximately 4.5 log_10_ CFU in the lungs. Mice were infected with approximately 4.5 log10 CFU. After infection, mice were randomized into treatment groups (12 mice per group). Untreated mice were sacrificed at the initiation of treatment to determine pretreatment CFU counts. Mice were initiated on treatment 3 d post-infection with one of four different regimens. The vehicle control group received a 200 μL subcutaneous injection of 20 mg/mL SHA-E_2_ without any drug. The second cohort of mice received a single 200 μL subcutaneous injection containing 600 μg of GFB (MedChemExpress). An additional cohort of mice received 10 doses of 60 μg of GFB administered by oral gavage once per day over the course of 14 days (5 doses for every 7 d). The experimental group received a 200 μL subcutaneous injection of 20 mg/mL SHA-E_2_ loaded with 600 μg of GFB. Lung CFU counts were assessed after 1 and 2 weeks of treatment by performing quantitative cultures of lung homogenates on OADC-enriched 7H11 agar (Difco Laboratories).

### Mouse Type 1 Diabetes Model

For *in vivo* studies with insulin, male C57BL/6J mice aged 6 to 8 weeks were purchased from Charles River Laboratories. After one week of acclimatation, mice were treated with 50 mg/kg of streptozotocin (STZ) (MilliporeSigma) for five consecutive days. STZ was dissolved at a concentration of 7.5 mg/ml in pH 4.5 sodium citrate buffer immediately before injection. Then, the mice’s blood glucose (BG) levels and weights were monitored. For blood glucose measurement, a drop of blood was collected from the tail and tested using a OneTouch UltraMini glucometer (LifeScan, Malvern, PA). Mice with blood glucose levels that exceeded 350 mg/dL for several sequential days were deemed diabetic and suitable for inclusion in the study.

Insulin-PBA was prepared from commercially purchased Insulin (35 µg/IU) as described in the chemical synthesis section of the methods. SHA-E_2_ hydrogels were prepared with Insulin-PBA with minor modifications to the protocol previously described. Insulin-PBA was dissolved at 8.4 mg/mL for 12 and 6 IU gels and 4.2 mg/mL for 3 IU gels in 2X PBS. PBS was used instead of HBSS to avoid injecting diabetic mice with glucose, which is part of the HBSS buffer. These solutions were then mixed 1:1 with a stock solution of 20 mg/mL of SHA-E_2_ dissolved in MQ water to form Insulin-PBA loaded hydrogels. Diabetic mice were then injected with 100 µL of 12 IU gels, 50 µL of 6 and 3 IU gels or with 3 IU of unmodified Insulin dissolved in 50 µL of 1X PBS. Right after treatment administration, blood glucose levels were measured every hour until 4 hours and every 2 hours until 10 hours. Blood glucose levels for mice that received insulin-PBA gel injections were subsequently monitored daily for 9 d. Mice that had blood glucose levels below 240 mg/dL were considered normoglycemic.

#### Statistical Analysis

Multiple group comparisons were calculated by one- or two-way ANOVA with Tukey’s multiple comparisons test. Statistical calculations were performed in GraphPad Prism 10. Statistical significance is denoted with asterisks as follows: *p <0.05; **p <0.01; ***p <0.001; ****p<0.0001. Error bars and reported error represents standard error of the mean (SEM) unless otherwise specified as standard deviation (SD) or as a 95% confidence interval.

