## Extended Data for "Enhanced dynamic covalent chemistry for the controlled release of small molecules and biologics from a nanofibrous peptide hydrogel platform"

Contents

Extended Data Fig. 1. Design and Development of SABER peptides. 2

Extended Data Fig. 2. *In vitro* release and drug stability. 3

Extended Data Fig. 3. *In vivo* release of Bortezomib (BTZ). 4

Extended Data Fig. 4. H&E and mass spectrometry imaging of injection site tissues. 5

Extended Data Fig. 5. Pharmacokinetics of GFB in 50 µL of 10 mg/mL Hydrogels. 6

Extended Data Fig. 6. *In vitro* characterization of SHA-$E_2$ and *in vivo* release of GFB. 7

Extended Data Fig. 7. Local release of PBA modified IgG. 8

Extended Data Fig. 8. Basal insulin delivery from SABER hydrogels. 9

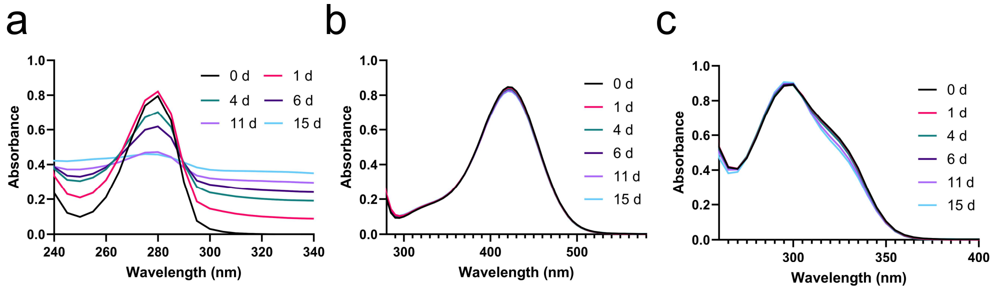


d

| | $K_{ARS}$ ($M^{-1}$) | $K_{DOPA}$ ($M^{-1}$) | $K_{nDOPA}$ ($M^{-1}$) | $K_{SHA}$ ($M^{-1}$) |
|---|---|---|---|---|
| **Tavaborole** | 1,030 ± 37.5 | 2,500 ± 149 | 1,420 ± 69.2 | 20,100 ± 829 |
| **Ixazomib** | --- | 1,920 ± 119 | 10,800 ± 3,230 | 17,600 ± 2,340 |
| **Ganfeborole (GFB)** | 1,220 ± 215 | 680 ± 70.3 | 141 ± 38.3 | 6,530 ± 747 |
| **3-Aminophenylboronic Acid** | 6,290 ± 121 | 6,000 ± 245 | 2,670 ± 426 | 38,800 ± 2,460 |

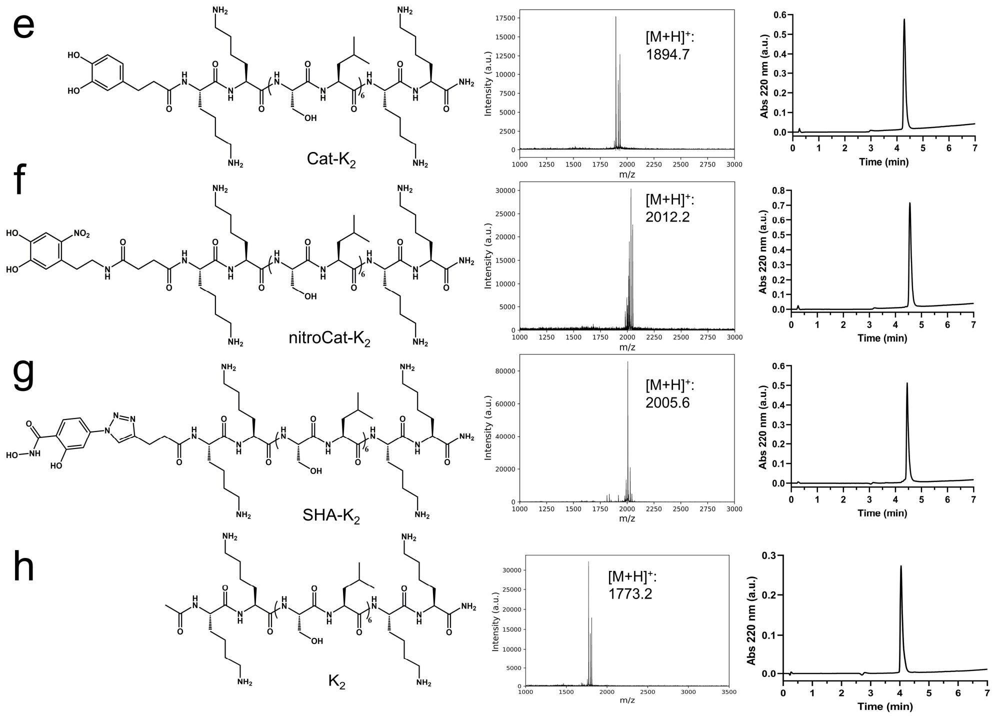


**Extended Data Fig. 1. Design and Development of SABER peptides.** a) Catechol oxidation and degradation occurred rapidly, appearing as changes in the UV-Vis spectrum of dopamine (DOPA) over time in pH 7.4 PBS. The increase in the baseline absorbance indicates increased turbidity of the sample as insoluble oxidation products form black precipitates in solution. b) The UV-Vis spectrum of 4-nitrocatechol (nDOPA) shows that the molecule is stable over the course of 15 days. c) Longitudinal analysis of the UV-Vis spectrum salicylhydroxamic acid (SHA) demonstrates that the molecule is largely stable with minor changes in absorbance observed on day 11 and day 15. d) Boronate ester equilibrium constants between chemically distinct boronic acids (BAs) and the dynamic covalent association motifs alizarin red S (ARS), nDOPA, SHA, and DOPA. For all BAs, SHA forms the strongest boronate ester interactions. Equilibrium constants are presented as the mean value of three replicates ± 1 SD. e-f) Chemical structures, mass spectra, and UPLC chromatograms of e) Cat-$K_2$, f) nitroCat-$K_2$, g) SHA-$K_2$, and h) $K_2$.

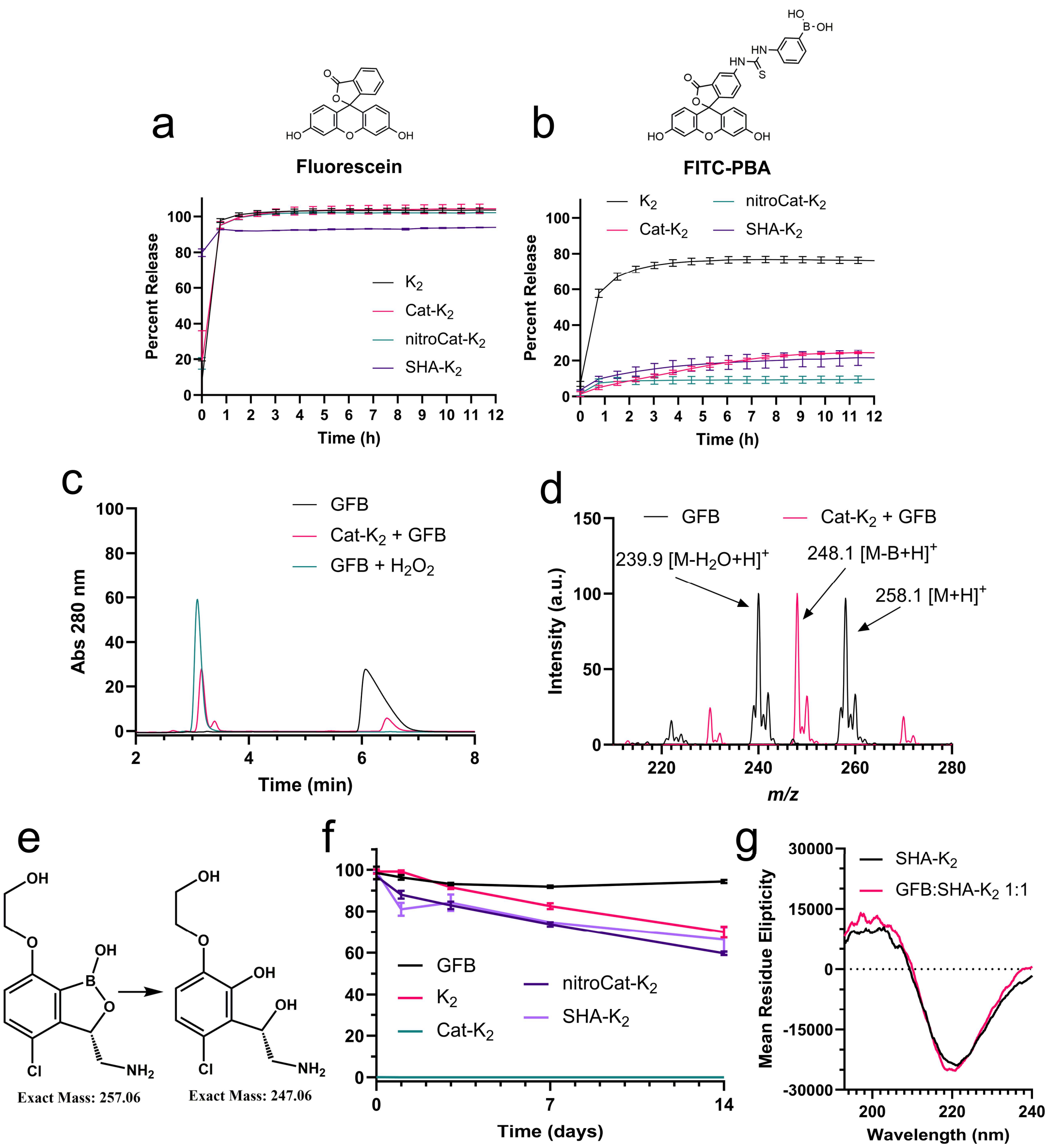


**Extended Data Fig. 2.** ***In vitro* release and drug stability.** a & b) Equilibrium fluorescence release results show that (a) unmodified fluorescein is rapidly released from all hydrogels, but (b) upon modification with phenylboronic acid (FITC-PBA), SABER hydrogels significantly prolong the release of the compound. Data is the mean (n=3) ± 1 SD. c) The change in UPLC retention time of GFB alone and GFB released from Cat-$K_2$ in UPLC suggests that the peptide may be degrading the drug. The new peak at 3.2 min seen in Cat-$K_2$ + GFB is also observed when the drug is reacted with 280 mM hydrogen peroxide, suggesting that this new peak corresponds to oxidation of GFB. d) Electrospray ionization mass spectrometry of GFB alone contains the expected mass of the drug at 258.1 *m/z* and an additional peak potentially due to a neutral water elimination from in-source fragmentation of the boronic acid. GFB released from Cat-$K_2$ has a peak at 248.1 *m/z*, (e) corresponding to a loss of a boron atom due to oxidative deboronation. f) Stability of GFB loaded in $K_2$, Cat-$K_2$, nitroCat-$K_2$, and SHA-$K_2$ quantified by UPLC shows that the majority of the drug remains stable over the course of 2 weeks in all hydrogels except for Cat-$K_2$, which immediately degrades the compound. All data is represented as the mean (n=3) ± 1 SD. g) SHA-$K_2$ without drug and the hydrogel maximally loaded (1:1 drug-to-peptide molar ratio) have the same β-sheet secondary structure as determined by a minimum at 220 nm seen by circular dichroism, suggesting that drug loading does not perturb peptide self-assembly.

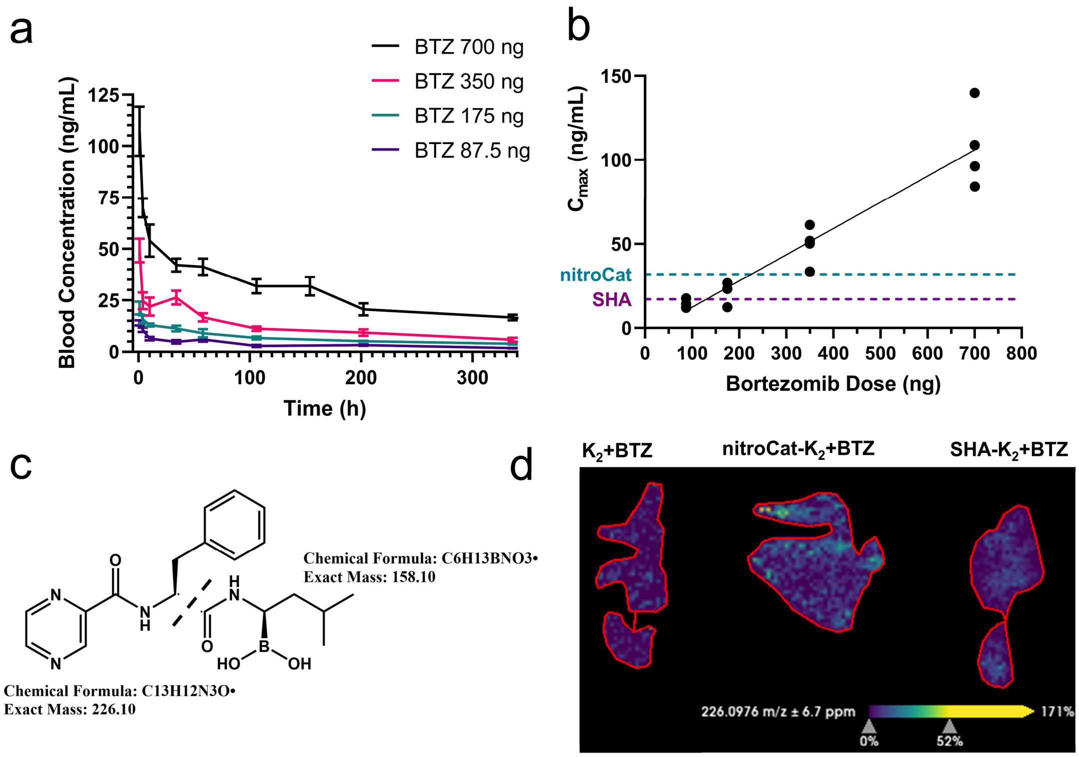


**Extended Data Fig. 3. *In vivo* release of Bortezomib (BTZ).** a) Pharmacokinetics of decreasing BTZ doses administered as subcutaneous boluses without hydrogel. Data is presented as the mean (n=4) ± SEM. b) The maximum circulating concentration ($C_{max}$) of BTZ decreases linearly with the initial dose. Dotted lines represent the $C_{max}$ for 700 ng of BTZ delivered from nitroCat-$K_2$ (blue) and SHA-$K_2$ (purple), indicating that a bolus BTZ dose of 175 ng yields the same $C_{max}$ as these hydrogel formulations loaded with 5-fold more drug. c) Chemical structure of the BTZ fragment observed in mass spectrometry imaging. d) $K_2$, nitroCat-$K_2$, and SHA-$K_2$ hydrogels loaded with 700 ng of BTZ imaged by mass spectrometry imaging *in vitro* show very little BTZ signal, suggesting that MDP peptides suppress the ionization of BTZ within the gels. This ionization suppression results in dark spots in mass spectrometry images at the location of the hydrogels.

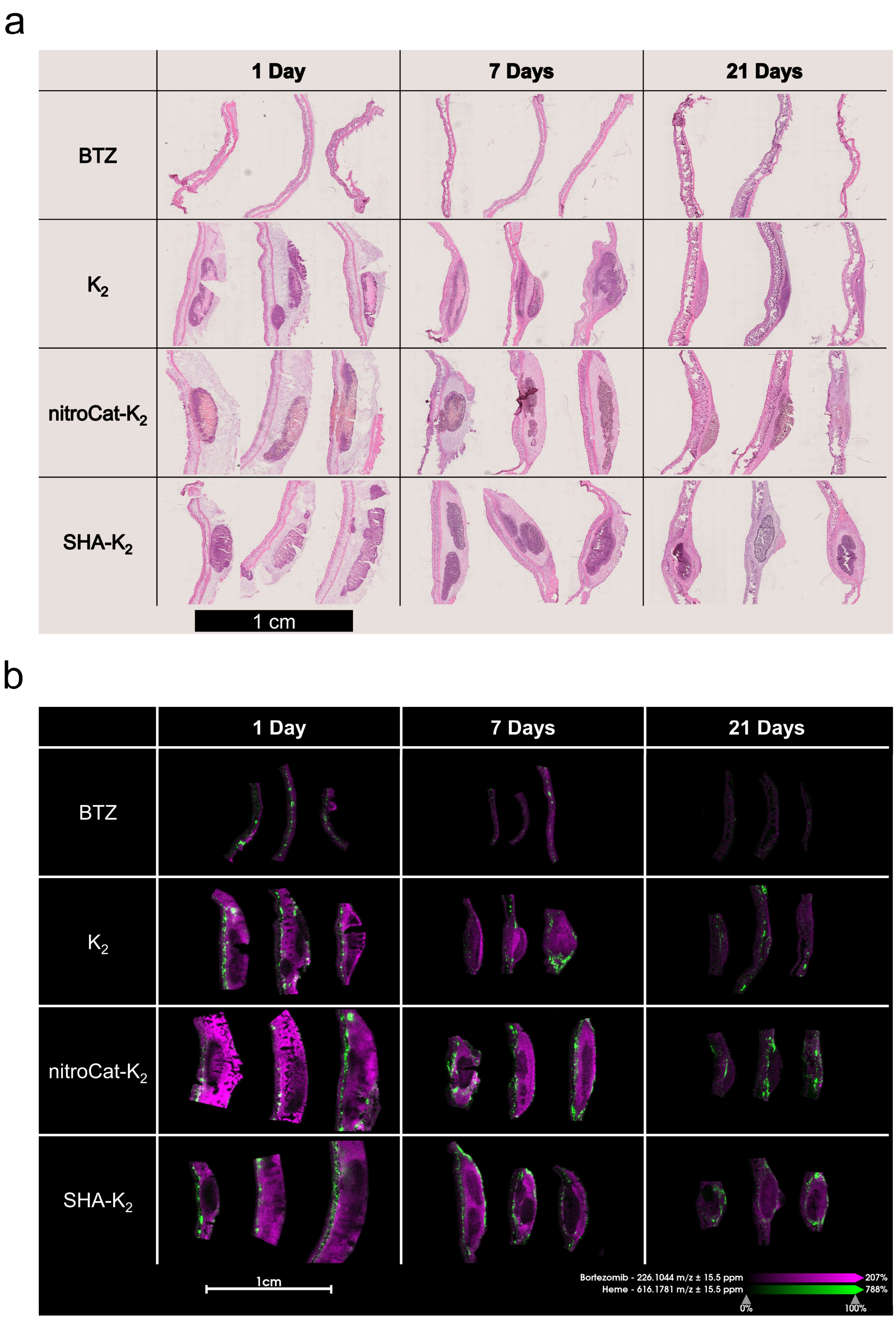


**Extended Data Fig. 4. H&E and mass spectrometry imaging of injection site tissues.** a) Tissue sections from mice that received subcutaneous injections of bortezomib (BTZ) alone or in a hydrogel at 1-, 7-, and 21-days stained with H&E. Large dark purple sections in $K_2$, nitroCat-$K_2$, and SHA-$K_2$ are the hydrogels in the skin samples. b) Mass spectrometry imaging of the same tissue samples stained with H&E shows that BTZ signal does not significantly overlap with that from heme (616.178 *m/z*), illustrating that the drug observed in the tissue is not in circulation but in the local environment of the injection site.

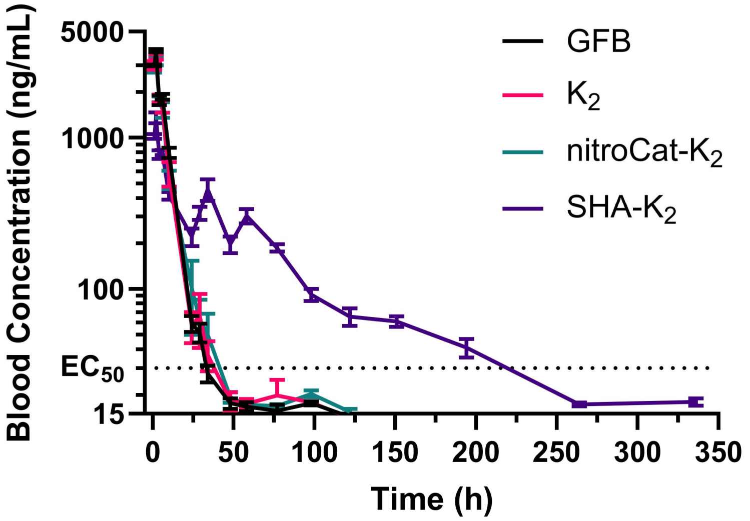


**Extended Data Fig. 5. Pharmacokinetics of GFB in 50 µL of 10 mg/mL Hydrogels.** Longitudinal concentrations of GFB in the blood of mice after a single subcutaneous injection of 75 µg of the drug loaded into 50 µL hydrogels or PBS (n=4). Injections of GFB without a hydrogel, GFB in $K_2$, and GFB in nitroCat-$K_2$ all led to the rapid release of the drug. SHA-$K_2$ hydrogels were able to retain GFB concentrations over the $EC_{50}$ for more than 200 h.

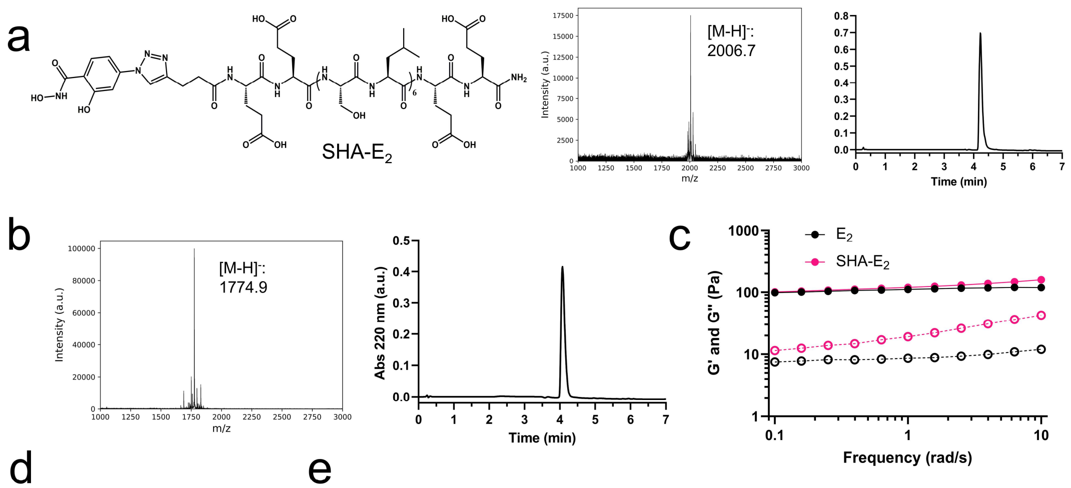


d

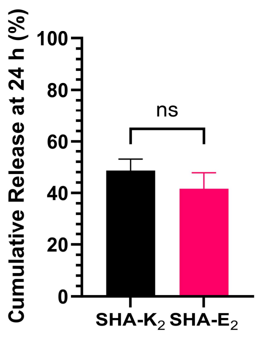


e

| | AUC (μg*h/mL) | $t_{1/2}$ (h) | $C_{max}$ (μg/mL) |
|---|---|---|---|
| **SHA-E$_2$ + 600 μg GFB** | 115 ± 9.22 | 154 ± 35.3 | 1.82 ± 0.20 |
| **Fold Change Due to Gel** | 1.33 | 10.2 | 0.097 |

**Extended Data Fig. 6. *In vitro* characterization of SHA-E$_2$ and *in vivo* release of GFB.** a) Chemical structure of SHA-E$_2$ with the mass spectrum and UPLC chromatogram of the material confirming the identity and purity of the peptide. B) Mass spectrum and UPLC chromatogram of the unmodified E$_2$ peptide. c) Frequency sweep collected by oscillatory rheology shows that SHA-E$_2$ hydrogels are more frequency dependent than unmodified E$_2$ and form slightly weaker gels as indicated by the reduced distance between the storage (G') and the loss (G") moduli. d) Cumulative release of GFB after 24 h from SHA-K$_2$ and SHA-E$_2$ are statistically similar, suggesting that changing the peptide used in SABER hydrogels does not compromise its ability to control the release of BA-containing small molecules. Data is presented as the mean of n=3 replicates ± 1 SD. e) Pharmacokinetic parameters extracted by performing a non-compartmental analysis on the *in vivo* release of GFB from SHA-E$_2$ show that using the SABER hydrogel improves drug exposure (AUC), half-life ($t_{1/2}$) and reduces the maximum circulating concentration ($C_{max}$) of the compound. Pharmacokinetic parameters are presented as the mean of n=4-5 replicates ± 1 SD.

a

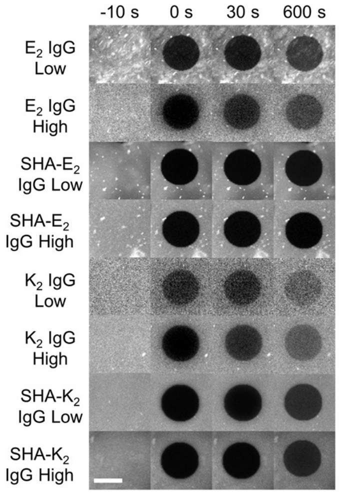


b

| Group | $t_{1/2}$ (min) | $M_f$ | $R^2$ |
|---|---|---|---|
| $E_2$ + IgG Low | 0.62 ± 0.04 | 0.37 ± 0.01 | 0.97 |
| SHA-$E_2$ + IgG Low | 2.45 ± 0.22 | 0.09 ± 0.07 | 0.96 |
| $E_2$ + IgG High | 0.64 ± 0.03 | 0.35 ± 0.01 | 0.97 |
| SHA-$E_2$ + IgG High* | 0.43 ± 0.04 | 0.05 ± 0.01 | 0.85 |
| $K_2$ + IgG Low | 0.63 ± 0.03 | 0.59 ± 0.01 | 0.97 |
| SHA-$K_2$ + IgG Low | 2.73 ± 0.12 | 0.25 ± 0.01 | 0.99 |
| $K_2$ + IgG High | 0.74 ± 0.04 | 0.47 ± 0.01 | 0.97 |
| SHA-$K_2$ + IgG High | 3.33 ± 0.19 | 0.23 ± 0.01 | 0.99 |

c

| Group | Percent Burst Release (Y0) | $t_{1/2}$ (days) | $R^2$ |
|---|---|---|---|
| IgG Low | 15.7 ± 5.1 | 0.4 ± 0.1 | 0.98 |
| IgG High | 0 ± 5.0 | 0.5 ± 0.1 | 0.99 |
| $E_2$ + IgG Low | 4.8 ± 7.2 | 0.7 ± 0.1 | 0.97 |
| $E_2$ + IgG High | 11.2 ± 7.4 | 0.8 ± 0.2 | 0.97 |
| SHA-$E_2$ + Low PBA | 21.8 ± 3.0 | 9.5 ± 0.7 | 0.99 |
| SHA-$E_2$ + IgG High | 20.4 ± 3.1 | 13.0 ± 0.9 | 0.98 |
| $K_2$ + IgG Low | 21.1 ± 6.3 | 2.2 ± 0.3 | 0.97 |
| $K_2$ + IgG High | 19.2 ± 5.8 | 4.1 ± 0.6 | 0.97 |
| SHA-$K_2$ + IgG Low | 22.1 ± 4.0 | 5.9 ± 2.4 | 0.98 |
| SHA-$K_2$ + IgG High | 22.5 ± 3.3 | 5.1 ± 0.6 | 0.99 |

**Extended Data Fig. 7. Local release of PBA-modified IgG.** a) Representative confocal images illustrating fluorescence recovery and photobleaching (FRAP) experiments. Fluorescence recovery was quantified by monitoring the return of fluorescence signal in the bleached region over 10 min using a 640 nm excitation laser. The white scale bar in the bottom left represents 40 µm. b) FRAP data (n=3 for each group) was fit to a first-order exponential equation to extract the FRAP half-time ($t_{1/2}$) and the mobile fraction ($M_f$). Loading PBA-modified IgG in SABER hydrogels reduced the $M_f$ and increased the $t_{1/2}$, suggesting that the rate of payload diffusion in these samples is slower. We observed minimal differences between IgG labeled with 11.4 PBAs per antibody (IgG high) and 2.4 PBAs per antibody (IgG low), demonstrating that the degree of labeling may not play a large role in controlling the rate of diffusion. All data were well-modeled by the first-order exponential equation and had $R^2$ values above 0.95 except for SHA-$E_2$ + IgG high (denoted with an asterisk), which was poorly fit by this model and thus extracted parameters may not accurately describe the data. c) *In vivo* release data of IgG with low and high degrees of PBA modeling was modeled with a first-order exponential equation to determine the half-life ($t_{1/2}$) and burst release from the site of injection. All SABER hydrogels had a moderate burst release of 20% but significantly extended the $t_{1/2}$ of the antibody at the injection site. All fits adequately modeled the data ($R^2$ > 0.95). All numerical data in this figure is presented as the mean (n=4) ± 95% confidence interval.

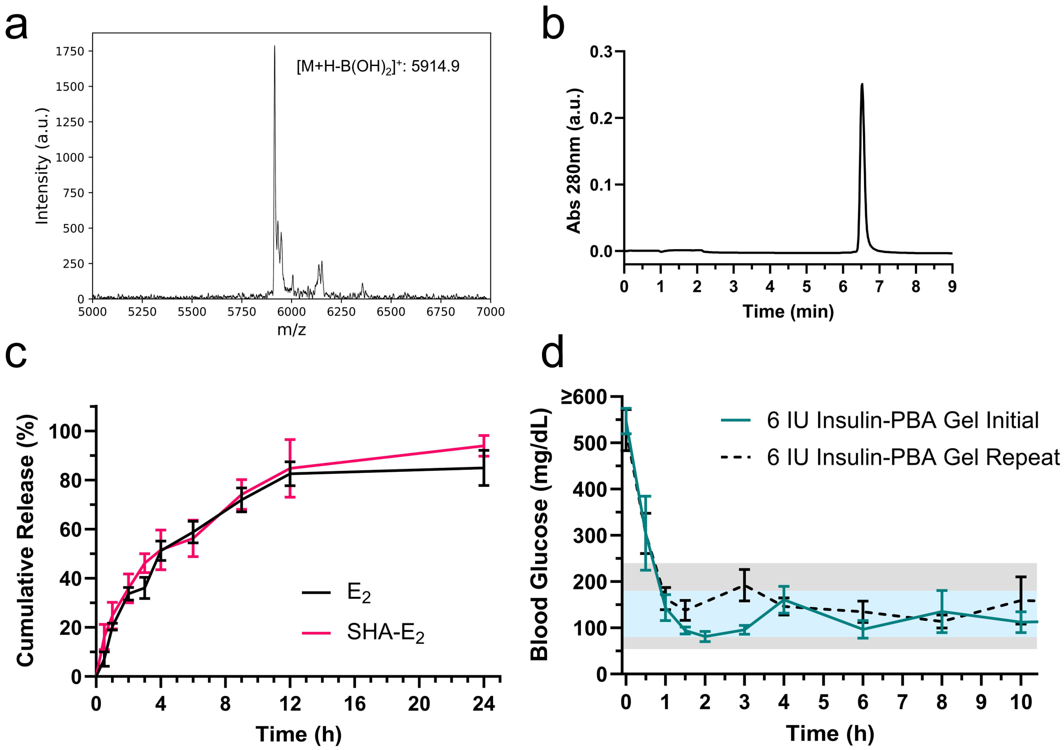


**Extended Data Fig. 8. Basal insulin delivery from SABER hydrogels.** a) Mass spectrum of PBA-modified insulin (insulin-PBA) showing that a single PBA was added to Ins. b) The UPLC chromatogram of the synthesized insulin-PBA confirmed the purity of the material. c) *In vitro* release of unmodified insulin from SHA-$E_2$ and $E_2$ illustrates that the PBA modification is necessary for the SABER peptide to delay the release of the payload. Data presented as the mean (n=3) ± 1 SD. d) First 10 h of the initial treatment of diabetic mice with 6 IU of insulin-PBA in SHA-$E_2$ plotted with a repeat treatment with the same formulation in the same mice 6 weeks later. The repeat dose resulted in statistically similar blood glucose levels to the initial dose at all time points. Data points for blood glucose measurements are the mean (n=5) ± SEM.
