## Supplementary Information for "Enhanced dynamic covalent chemistry for the controlled release of small molecules and biologics from a nanofibrous peptide hydrogel platform"

**Table of Contents**

Table S1. UPLC methods for quantification of *in vitro* drug release. SI-2

Table S2. Bortezomib and apatinib MRM ion transitions used to quantify blood concentrations by QqQ LC-MS. SI-2

Table S3. Ganfeborole and apatinib MRM ion transitions used to quantify blood concentrations by QqQ LC-MS. SI-2

Drug Loading Discussion SI-3

Fluorescence Recovery After Photobleaching Discussion SI-3

Supplementary Methods SI-4

- Chemical Synthesis SI-4
- UV-Vis Oxidation Study SI-10
- Boronate Ester Association Constant Determination SI-11
- *In Vitro* TLR7 Activity Assay SI-12
- Fluorescence Recovery After Photobleaching SI-12

Supplementary Information References SI-13

**Table S1.** UPLC methods for quantification of *in vitro* drug release.

| | Column | Column Temp (°C) | Solvent A | Solvent B | Gradient | Wavelength (nm) |
|---|---|---|---|---|---|---|
| Bortezomib | C18 | 25 | Water + 0.05% TFA | Acetonitrile + 0.05% TFA | 15-35% B at 5% B/min | 270 |
| Ixazomib | C18 | 25 | Water + 0.05% TFA | Acetonitrile + 0.05% TFA | 25%-65% B at 10% B/min | 210/280 |
| Tavaborole | C18 | 25 | Water + 0.05% TFA | Acetonitrile + 0.05% TFA | Isocratic 10% B for 7.5 min | 210/265 |
| 1V209 & 1V209-PBA | C18 | 25 | Water + 0.05% TFA | Acetonitrile + 0.05% TFA | 5-28% B at 2.875% B/min | 220/280 |
| Ganfeborole | C18 | 45 | Water + 0.05% TFA | Methanol + 0.05% TFA | Isocratic 12% B for 7.5 min | 230/280 |
| Insulin & Insulin-PBA | C4 | 25 | Water + 0.05% TFA | Acetonitrile + 0.05% TFA | 25-45% B at 5% B/min | 280 |

**Table S2.** Bortezomib and apatinib MRM ion transitions used to quantify blood concentrations by QqQ LC-MS.

| Compound Name | Precursor Ion (Da) | Product Ion (Da) | Fragmentor Voltage (V) | Collision Energy (eV) | Note |
|---|---|---|---|---|---|
| Apatinib | 398.2 | 92 | 152 | 46 | Qualifier |
| Apatinib | 398.2 | 80.1 | 152 | 62 | Qualifier |
| Apatinib | 398.2 | 212 | 152 | 30 | Primary |
| Brotezomib | 367.2 | 226 | 136 | 19 | Primary |
| Brotezomib | 367.2 | 208 | 136 | 31 | Qualifier |
| Brotezomib | 367.2 | 79.1 | 136 | 54 | Qualifier |

**Table S3.** Ganfeborole and apatinib MRM ion transitions used to quantify blood concentrations by QqQ LC-MS.

| Compound Name | Precursor Ion (Da) | Product Ion (Da) | Fragmentor Voltage (V) | Collision Energy (eV) | Note |
|---|---|---|---|---|---|
| Apatinib | 398.2 | 92 | 152 | 46 | Qualifier |
| Apatinib | 398.2 | 212 | 152 | 30 | Primary |
| Ganfeborole | 258.1 | 222 | 106 | 13 | Primary |
| Ganfeborole | 258.1 | 187 | 106 | 25 | Qualifier |

### Drug Loading Discussion

Since each BACSM molecule must bond to a diol-containing peptide to avoid release via diffusion, we posited that increasing the ratio of available SHA motifs to BACSMs would increase the likelihood of boronate ester reformation after hydrolysis and thereby increase drug retention. To investigate how a surplus of peptide-displayed SHA motifs might modify drug release kinetics, we varied the molar ratio of drug-to-peptide in SHA-$K_2$ gels. $K_2$ and SHA-$K_2$ hydrogels were prepared at different levels of drug loading ranging from a 1:1 to 1:10 drug-to-peptide molar ratio while keeping the peptide concentration constant at 10 mg/mL (~5 mM). The release of GFB from $K_2$ was fast, as expected, and did not significantly change as a function of drug loading over this range (Fig. 2e). In contrast, the release of drug from SHA-$K_2$ was sensitive to drug loading (Fig. 2e). After 24 h, the 1:1 drug-to-peptide SHA-$K_2$ gel loaded with the most drug released 45.4 ± 1.1%, significantly more than the 1:2 (35.1 ± 0.6%), 1:4 (31.5 ± 0.6%), and 1:10 (31.3 ± 0.7%) gels (Fig. 2f). There were no statistical differences between the three lower loadings, suggesting that drug loading has a negligible impact on release rate once the number of SHA motifs significantly exceeds the number of drug binding partners. We also confirmed that the highest degree of drug loading did not perturb self-assembly by CD analysis of the 1:1 SHA-$K_2$ hydrogel (Extended Data Fig. 2g). SHA-$K_2$ significantly delayed the release of GFB compared to unmodified $K_2$ at all molar ratios tested, which indicates that these hydrogels can be “fully” loaded with one BACSM per dynamic covalent attachment motif and still function effectively as a drug delivery vehicle.

### Fluorescence Recovery After Photobleaching Discussion

We then investigated diffusion of PBA-labeled IgG in SHA-$K_2$ and SHA-$E_2$ hydrogels using fluorescence recovery after photobleaching (FRAP) studies. After the fluorescence signal in three distinct locations in the gel was photobleached (Extended Data Fig. 7a), the diffusion of IgG into the bleached area over time was significantly higher in unmodified MDPs compared to SHA-functionalized peptides (Fig. 5b & c). We observed a 50% decrease in mobile fraction of IgG, and an over 100% increase in the time it takes to recover half the final fluorescence intensity ($t_{1/2}$) in SHA-$K_2$ compared to $K_2$ (Extended Data Fig. 7b). The decrease in the rate of IgG diffusion within SHA-modified gels was further enhanced in SHA- $E_2$. The mobile fraction of IgG in SHA-$E_2$ was less than 25% of the mobile fraction of IgG observed in unmodified $E_2$ (Extended Data Fig. 7b). Interestingly, the degree of PBA labeling did not have a meaningful impact on the FRAP data, as high and low degrees of IgG labeling displayed very similar results despite the nearly 5-fold difference in the degree of PBA labeling.

### Supplementary Methods

#### Chemical Synthesis

All chemicals, unless otherwise specified, were purchased from Fisher Scientific (Pittsburgh, PA) or MilliporeSigma (Burlington, MA). Flash chromatography was performed on silica gel (SiliaFlash P60).

##### Synthesis of 4-nitrodopamine

Dopamine (500 mg, 2.6 mmol) and sodium nitrate (630 mg, 9.1 mmol) were dissolved in 15 mL of water and cooled in an ice bath. Once cold, the solution was stirred vigorously and 2.5 mL of 20% sulfuric acid was added dropwise, which resulted in a yellow precipitate that was collected by vacuum filtration. The product was washed 3X with cold water and 3X with cold methanol and dried under vacuum with a 50% yield. $^1$HNMR (600HZ, DMF-d7) δ = 3.35 (m, 4H), 7.27 (s, 1H), 7.53 (s, 1H). ESI-MS expected for $C_8H_{11}N_2O_4^+$ [M+H]$^+$ 199.2, observed [M+H]$^+$ 199.1.

Synthesis of azido-salicylhydroxamate-O$^t$Bu **2**

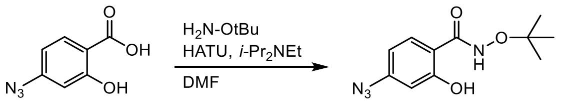


Azido-salicylic acid **1** was synthesized as previously reported.[1] Solid 4-azidosalicylic acid **1** (2.5 g, 14 mmol) and Hexafluorophosphate azabenzotriazole tetramethyl uronium (HATU) (5.3 g, 14 mmol) were dissolved in dry DMF (50 mL) under nitrogen. DIEA (6.3 mL, 36 mmol) was added at 0 °C and the solution was stirred at rt for 10 min. Solid *O*-(tert-Butyl)hydroxylamine hydrochloride (1.96 g, 15 mmol) was added at 0 °C and the solution was stirred at rt for 64 h. The solution was diluted with EtOAc (200 mL) and washed with sat. $KHSO_4$ (3 x 120 mL), 1:1 $H_2O$/brine (3 x 120 mL), and brine (200 mL). The organic layer was dried over $MgSO_4$, filtered, and solvent was removed under vacuum. The residue was purified by flash column (25% EtOAc/Hexane) to afford **2** as a white powder (1.29 g, 37%). $^1$H NMR (600 MHz, DMSO-d6): δ 7.76 (d, *J* = 8.50 Hz, 1H), 6.67 (m, 1H), 6.62 (s, 1H), 1.24 (s, 9H) $^{13}$C NMR (151 MHz, DMSO-d6): δ 167.6, 160.7, 144.8, 132.6, 130.3, 110.4, 107.4, 82.1, 26.8 ESI-MS expected for $C_{11}H_{14}N_4O_3$ $[M+H]^+$: 251.3, found 251.1.

Synthesis of COOH-SHA-OtBu **3**

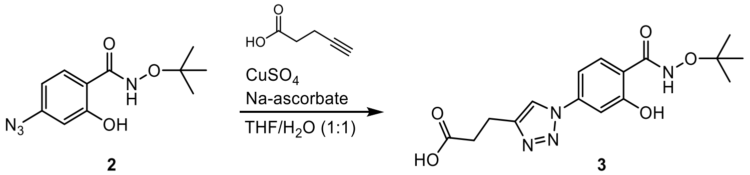


Solid azido-salicylhydroxamate-OtBu **2** (880 mg, 3.5 mmol) and 4-pentynoic acid (380 mg, 3.9 mmol) were dissolved in THF (4.4 mL). Sodium ascorbate (850 mg, 4.2 mmol) in $H_2O$ (2.2 ml) was added followed by copper(II) sulfate pentahydrate (270 mg, 1.1 mmol) in $H_2O$ (2.2 mL). The solution was stirred vigorously at rt for 2 h. Solvent was removed under vacuum and the residue was suspended in MeOH, filtered, and the supernatant was concentrated under reduced pressure and then purified by flash column (20% MeOH/$CH_2Cl_2$) to afford **3** as an off-white powder (1.08 g, 88%). $^{1}$H NMR (600 MHz, DMSO-d6): δ 8.66 (s, 1H), 7.89 (d, *J* = 8.56 Hz, 1H), 7.47 (s, 1H), 7.42 (d, *J* = 8.53 Hz, 1H), 2.93 (t, *J* = 7.47 Hz, 2H), 2.67 (t, *J* = 7.51 Hz, 2H), 1.26 (s, 9H) $^{13}$C NMR (151 MHz, DMSO-d6): δ 174, 167.0, 160.1, 147.7, 140.3, 130.4, 120.8, 115.6, 110.2, 107.9, 82.2, 33.2, 26.9, 21.1 ESI-MS expected for $C_{16}H_{20}N_4O_5$ $[M+H]^+$: 349.4, found 349.1.

<u>Synthesis of 3-(3-(3',6'-dihydroxy-3-oxo-3H-spiro[isobenzofuran-1,9'-xanthene]-5-yl)thioureido)phenylboronic acid (FITC-PBA)</u>

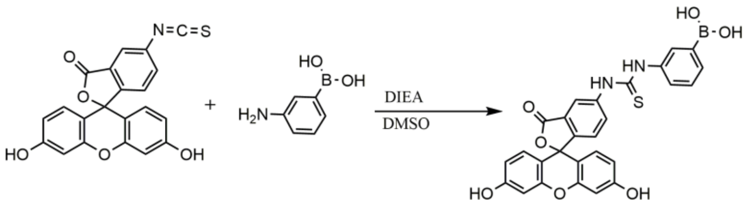


Solid fluorescein isothiocyanate (FITC) (100 mg, 0.257 mmol) and 3-aminophenylboronic acid (Ambeed, Arlington Heights, IL) (52.8 mg, 0.386 mmol) were dissolved in 0.5 mL in DMF. Diisopropylethylamine (DIEA) (44.8 μL, 0.257 mmol) was added to the reaction under agitation. The reaction was allowed to proceed overnight protected from light. Excess methanol (10 mL) was added to the reaction and the solvent was removed in vacuo. The product was redissolved in a minimal volume of methanol and the product was precipitated into chloroform. The solid was isolated by vacuum filtration and dried under vacuum for a final yield of 38%. ESI-MS expected for $C_{27}H_{18}BN_2O_7S^-$ $[M-H]^-$ 525.3, observed $[M-H]^-$ 525.21.

Synthesis of (4-((4-((6-amino-2-(2-methoxyethoxy)-8-oxo-7,8-dihydro-9H-purin-9-yl)methyl)benzamido)methyl)phenyl) boronic acid (1V209-PBA)

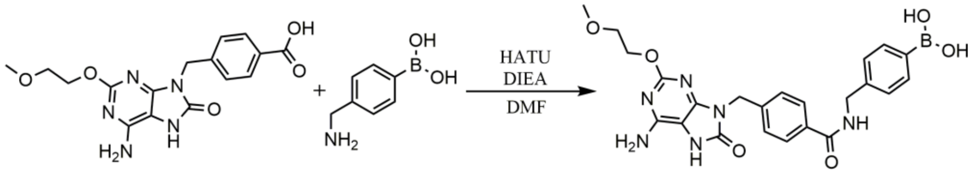


Solid 4-((6-amino-2-(2-methoxyethoxy)-8-oxo-7,8-dihydro-9H-purin-9-yl)methyl)benzoic acid (1V209) (Ambeed) (50 mg, 0.139 mmol) and HATU (P3 biosystems, Louisville, KY) (55.6 mg, 0.146 mmol) were dissolved in a minimal volume of DMF and stirred. To the solution, 144.5 uL (0.835 mmol) of DIEA was added and the reaction was allowed to stir for two min. Afterward, 52.2 mg (0.278 mmol) of 4-aminomethylphenylboronic acid (Ambeed) was added and the reaction was allowed to proceed at ambient temperature for 2 h. The reaction mixture was then purified by high-performance liquid chromatography (Shimadzu Corp., Kyoto, Japan) on an XBridge BEH C18 column (Waters Corp., Milford, MA) using a solvent system of water and acetonitrile with 0.05% TFA. The purified product was frozen and lyophilized for a final yield of 27%. $^{1}$H NMR (600 MHz, DMSO-d6): δ: 10.0 (s, 1H), 7.85 (d, J=7.8 Hz, 2H), 7.73 (d, J=7.8 Hz, 2H), 7.37 (d, J=8.4 Hz, 2H), 7.25 (d, J=7.8 Hz, 2H), 6.5 (s, 2H), 4.92 (s, 2H), 4.5 (d, J=6 Hz, 2H), 4.25 (t, J=4.38 Hz, 2H), 3.58 (t, J=4.86 Hz, 2H), 3.26 (s, 3H). ESI-MS expected for $C_{23}H_{26}BN_6O_6^+$ $[M+H]^+$ 493.2, observed $[M+H]^+$ 493.2.

Solid-Phase Peptide Synthesis

Peptides were synthesized manually or on an AAPPTec Focus XC autosynthesizer (AAPPTec, Louisville, KY) using standard fluorenylmethyloxycarbonyl (fmoc)-based chemistry. Fmoc-protected low-loading MBHA rink-amide and fmoc-protected amino acids were procured from Novabiochem (MilliporeSigma). Fmoc deprotection was accomplished with two steps of excess 25% v/v piperidine in DMF:DMSO 1:1 for 5 min each. Successful deprotection was verified using the ninhydrin test for primary amines. Coupling was achieved with the addition of preactivated Fmoc-protected amino acids (4 equiv.), HATU (3.99 eq.) (P3 biosystems) and DIEA (6 eq.) in a minimal amount of dimethylformamide (DMF): dimethyl sulfoxide (DMSO) 1:1 for 20-45 min at RT. The process was repeated until the peptide was complete. In the case of the $K_2$ and $E_2$ peptide N-terminal acetylation was performed with the addition of acetic anhydride (200 eq.) and DIEA (75 eq.) in dichloromethane (DCM) two times for 45 mins.

For all other SABER peptides, the N-terminus was modified with a boronic acid binding motif instead of being acetylated. Cat-$K_2$ and SHA-$K_2$/SHA-$E_2$ were generated by coupling 4 equiv. of 3-(2,2-Dimethylbenzo[d][1,3]dioxol-5-yl)propanoic acid or 2 equiv. of COOH-SHA-OtBu, respectively, using the same coupling protocol described above. NitroCat-$K_2$ was synthesized by reacting the N-terminus of the peptide with 2 equiv. of bis(2,5-dioxopyrrolidin-1-yl) succinate (Ambeed) and 4 equiv. of DIEA for 30 min in DMF/DMSO. Successful coupling was verified by using the ninhydrin test for primary amines. The resin was then washed 3X with DCM and DMF and then reacted with 4 equiv. of nitrodopamine and 10 equiv of DIEA in DMF/DMSO. The reaction was allowed to proceed overnight.

Peptides were cleaved from the resin by reaction with a cleavage cocktail composed of trifluoroacetic acid (TFA):Anisole:triisopropylsilane:ethylene-1,2-dithiol:$H_2O$ 90:2.5:2.5:2.5:2.5 for 3 h at RT. The t-butyl protecting group on hydroxamic acid of SHA-$K_2$ and SHA-$E_2$ was removed by heating the cleavage cocktail to 50 °C for an additional hour. TFA was then evaporated with a stream of nitrogen to a minimal volume ( ~ 1 mL) and peptide was precipitated with ice-cold diethyl ether.

Peptides were purified by high performance liquid chromatography (Shimadzu Corp., Kyoto, Japan) on an XBridge BEH OBD C4 column (Waters Corp.). Positively charged peptides were purified over a gradient of 5-50% acetonitrile with 0.05% TFA in water with 0.05% TFA. Negatively charged peptides were purified over a gradient of 5-35% acetonitrile with an ammonium acetate buffer (5 mM ammonium and 4 mM acetic acid) in water with the same buffer with a pH of 8.5. The purity of all peptides was verified by ultraperformance liquid chromatography

(UPLC) (Waters Corp.) and matrix-assisted laser desorption-ionization mass spectrometry (MALDI-MS) (Bruker Daltonics, Billerica, MA).

IgG PBA Labeling

Rabbit IgG (MilliporeSigma) was dissolved at 10 mg/mL (6.66 x $10^{-5}$ M) in pH 8.2 sodium bicarbonate buffer (0.2 M) and (4-(((2,5-Dioxopyrrolidin-1-yl)oxy)carbonyl)phenyl)boronic acid (Ambeed) was dissolved at 0.05 M in DMSO. To make the high degree of PBA-labeled IgG, 20 equivalents of (4-(((2,5-Dioxopyrrolidin-1-yl)oxy)carbonyl)phenyl)boronic acid was then added to the dissolved IgG while 5 equivalents was added for the low degree of labeling. The reaction was stirred at ambient temperature overnight. The product was then purified using a 0.5 mL size exclusion Spin-X centrifugal filter with a 50 kDa cutoff (Corning, Corning, NY). The degree of labeling was determined by quantifying the amount of unreacted (4-(((2,5-Dioxopyrrolidin-1-yl)oxy)carbonyl)phenyl)boronic acid that eluted out in the filtrate by UV-Vis. This characterization method was validated by an alizarin red S assay for the detection of BAs. These reactions generated IgG material with an average of 11.4 and 2.4 PBAs per IgG molecule, corresponding to an approximately a 50% PBA labeling efficiency. The IgG was then subsequently labeled with the NHS ester-functionalized AZDye 647 fluorophore (Vector Laboratories, Newark, NJ) using the protocol published by the supplier to achieve a degree of labeling of 0.2 fluorophores per IgG. The product was then purified by size exclusion using a 15 mL 50 kDa Amicon centrifugal filter (MilliporeSigma) against water and stored as a lyophilized powder.

Synthesis of insulin-PBA

Insulin labeled with a single phenylboronic acid on the B29 Lys residue (insulin-PBA) was prepared following previously published protocols.[2,3] In brief, 5.6 mg of (4-(((2,5-Dioxopyrrolidin-1-yl)oxy)carbonyl)phenyl)boronic acid (Ambeed) was dissolved in 1 mL of acetonitrile (21.12 mM; 1.2 equiv.) and 100 mg of recombinant human insulin (Thermo Fisher Scientific, Waltham, MA) was dissolved in 1 mL of 0.1 M sodium carbonate buffer set to pH 10.2 (17.6 mM; 1 equiv.). After complete dissolution of the insulin, the pH was adjusted back to pH 10.2 with NaOH. The two solutions were then mixed 1:1 and stirred for 1 h at ambient temperature. The reaction was then quenched through the addition of 88 μL of 0.2 M methylamine (MilliporeSigma). The pH of the reaction solution was then adjusted to pH 5.5 with 6 M HCl at which point the product crashed out of solution as a white precipitate. The solution was then cooled at 4 °C for 30 min and the precipitate was isolated by centrifugation at 10,000 rcf for 2 min. The product was then dissolved in 4 mL of DMSO with 0.1% TFA and purified by HPLC on an XSelect OBD BEH C18 column (Waters Corp.) using a solvent system of water and acetonitrile both with 0.5% TFA. The purity and identity of the product was assessed by UPLC and MALDI-MS using sinapic acid (MilliporeSigma) as the matrix. Expected $[M+H]^+$: 5957.9; observed $[M+H-B(OH)_2]^+$: 5914.9. The loss of 43 Da is consistent with in-source fragmentation of the boronic acid. The purified product was stored as a lyophilized solid until use.

**UV-Vis Oxidation Study**

Oxidation of dopamine, 4-nitrodopamine, and SHA were analyzed by qualitative changes in absorbance spectra from 200 to 800 nm with a Cary 60 UV Vis Spectrophotometer (Agilent Technologies, Santa Clara, CA) using a quartz cuvette. Compounds of interest were prepared in 1X PBS at concentrations such that the maximum absorbance of the sample would not exceed 1 absorbance unit. Samples were stored at ambient temperature in the dark at 25 °C for the duration of the experiment. Spectra were collected at 0, 1, 4, 6, 11, and 15 d after preparation to monitor the changes in the samples over time.

**Boronate Ester Association Constant Determination**

The determination of boronate ester equilibrium by competitive binding assay were determined by adapting previously described protocols.[4,5] To determine the BACSM-alizarin red S (ARS) binding constants ($K_{ARS}$) for aryl BACSMs, stock solutions of ARS and BACSM were prepared in 1X PBS and corrected to pH 7.4 as needed. Titrations of decreasing BACSM concentration (starting at 2 mM) in constant ARS concentration (9 µM) were prepared in triplicate. Fluorescence was measured (468/572 nm Ex/Em) with a Tecan Infinite M200 Pro Plate Reader (Zurich, Switzerland) using black 96-well plates. The ARS association constant ($K_{ARS}$) was determined using equations previously described.[4] $K_{ARS}$ was then used to determine the boronate ester equilibrium constant between the BACSM and dopamine (DOPA) and SHA by preparing serial dilutions in triplicate of these two compounds (starting at 5 mM) with constant concentrations of ARS (9 µM) and BACSM (2 mM). The fluorescence (468/572 nm Ex/Em) of these dilutions was measured using a microplate reader and $K_{DOPA}$ and $K_{SHA}$ were calculated as previously reported.[4]

The absorbance spectrum of 4-nitrodopamine (nDOPA) overlaps with the excitation wavelength of ARS; however, its $\lambda_{max}$ is sensitive to boronate ester formation (observed as a blue-shift from 420 nm). Thus, to determine the BACSM-nDOPA binding constants ($K_{nDOPA}$), serial dilutions of each BACSM (starting at 2 mM) were prepared in triplicate with a constant concentration of nDOPA (0.1 mM) in pH 7.4 1X PBS. The absorbance at 420 nm was measured by microplate reader using a quartz 96 well plate. $K_{nDOPA}$ was determined with the same equations used to calculate $K_{ARS}$, where 1-absorbance was substituted for fluorescence intensity. Since ARS fluorescence changes only occur with aryl BAs, $K_{nDOPA}$ was used to determine binding constants between Ixazomib (IXB) and DOPA ($K_{DOPA}$) or SHA ($K_{SHA}$). In brief, serial dilutions of DOPA or SHA (starting at 2 mM) were prepared in triplicate with constant IXB (2 mM) and nDOPA (0.1 mM). The absorbance at 420 nm of these solutions was measured and used to calculate $K_{DOPA}$

or $K_{SHA}$ using the previously calculated $K_{nDOPA}$ for IXB with 1-absorbance substituted for fluorescence intensity in the original equation previously published.[4]

#### *In Vitro* TLR7 Activity Assay

The ability of 1V209-PBA to activate TLR 7 was determined using the HEK-Blue mTLR7 reporter cell line (InvivoGen, San Diego, CA) using the protocol provided by the supplier. HEK Blue cells were plated at a density of $4.0 \times 10^4$ cells/well. The cells were exposed to 2-fold serial dilutions of 1V209 and 1V209-PBA over a concentration range of 20 – 0.02 µM with 2% DMSO and incubated at 37 °C and 5% $CO_2$ for 20-24 h. After incubation, 20 µL of supernatant was collected from each well and incubated with 180 µL of QUANTI-Blue detection media (InvivoGen) at 37 °C for 6h in a 96-well plate. TLR7 activation was quantified by reading the absorbance of the detection solutions at 630 nm using a microplate reader.

#### Fluorescence Recovery After Photobleaching (FRAP)

Hydrogels loaded with 2 mg/mL of AZ647-labeled IgG-PBA were prepared as previously described, and 10 uL of each hydrogel was pipetted onto a 25 x 75 x 1 mm Diamond White glass microscope slide (Globe Scientific Inc., Mahwah, NJ) with a secure seal imaging spacer with a 9 mm diameter and 0.12 mm depth (Electron Microscopy Sciences, Hatfield, PA). The sample in the spacer was then covered with 24 x 40 mm cover glass (Corning Inc.). FRAP experiments were conducted on a Nikon A1 Confocal microscope utilizing the NIS-Elements AR 5.21.03 software (Nikon, Tokyo, Japan) Galvano mode with an 20X objective lens following a previously published protocol.[58] Total field of view was 512 x 512 pixels. The field of view was focused until the fluorescence signal from the 640 nm laser line was maximized. Bleaching was performed on three 50 µm diameter spots at a scan speed of 1 frame per second using 486 nm, 561 nm, and 640 nm lasers at 100% intensity for 15.16 sec with a 10 min recovery imaging period using the 640 nm laser. Images were taken every 2 sec for the first min of recovery and then every 10 sec for the following 9 min. Three different bleach spots were measured and recorded for each sample. The fluorescence intensity at each time point during the recovery was normalized to the pre-bleached fluorescence intensity of the spot and the data was fit to a first-order exponential equation using a least-squares regression in Prism 10 to extract the fluorescence recovery half-time ($t_{1/2}$) and the mobile fraction ($M_f$) as previously described.[59] Error for all parameters were reported as 95% confidence intervals.
